# On-tissue spatially-resolved glycoproteomics guided by N-glycan imaging reveal global dysregulation of canine glioma glycoproteomic landscape

**DOI:** 10.1101/2020.10.02.324434

**Authors:** Stacy A. Malaker, Jusal Quanico, Antonella Raffo Romero, Firas Kobeissy, Soulaimane Aboulouard, Dominique Tierny, Carolyn R. Bertozzi, Isabelle Fournier, Michel Salzet

## Abstract

Here we present an approach to identify N-linked glycoproteins and deduce their spatial localization using a combination of MALDI mass spectrometry N-glycan imaging and spatially-resolved glycoproteomic strategies. We subjected formalin-fixed, paraffin-embedded glioma biopsies to on-tissue PNGaseF digestion and MALDI imaging and found that the glycan HexNAc4-Hex5-NeuAc2 was found to be predominantly expressed in necrotic regions of high-grade canine gliomas, whereas high mannose HexNAc2-Hex5 was predominantly found in benign regions. To determine the underlying sialo-glycoprotein, various regions in adjacent tissue sections were subjected to microdigestion and the extracts were analyzed by LC-MS/MS without further glycopeptide enrichment. Results identified haptoglobin, which is involved in iron scavenging that presents aberrant fucosylation/sialylation in various cancers, as the protein associated with HexNAc4-Hex5-NeuAc2. Additionally, we identified several high-mannose (Hex2-HexNAc5) glycopeptides enriched in benign regions. To the best of our knowledge, this is the first report that directly links glycan imaging with intact glycopeptide identification. In total, our spatially-resolved glycoproteomics technique identified over 400 N-glycosylated, O-GalNAcylated, O-mannosylated, and S- and O-GlcNAcylated glycopeptides from over 30 proteins, demonstrating the diverse array of glycosylation present on the tissue slides and the sensitivity of our technique. While N-glycosylation and O-mannosylation were similar between benign and tumor/necrotic sections, S- and O-GlcNAc glycopeptides were significantly deceased in tumor/necrotic sections, whereas sialylated O-GalNAc glycopeptides were significantly upregulated. Ultimately, this proof-of-principle work demonstrates the capability of spatially-resolved glycoproteomics to complement MALDI-imaging technologies in understanding dysregulated glycosylation in cancer.

## INTRODUCTION

Canine glioma is a spontaneous non-human intracranial neoplasm that has recently gained attention as a viable preclinical model for comparative oncology.^1–3^ Compared to neurosphere cultures and patient-derived homogeneous xenografts transplanted on immune-deficient mice, canine glioma offers a lesser “translational gap” with human glioma. Both human and canine gliomas are highly heterogeneous and present similar histopathological, immunological, molecular, and clinical characteristics. Additionally, the animals are immunocompetent and they exhibit similar responses to chemotherapy, radiotherapy, and stereotactic radiosurgery.^4^ Further, being household pets, dogs are exposed to similar environmental pressures as their owners and, like humans, are also susceptible to various spontaneous malignancies.^3^ Canine glioma can thus serve as an intermediate model that can be exploited for preclinical evaluation of new treatments, providing putative candidates that not only are expected to have better chances of passing phase II clinical trials, but also will be accessible to companion animals. Human glioma is a high-grade brain tumor with poor prognosis and currently, the only two treatment options are FDA-approved: temozolamide and bevacizumab.^5^ In an effort to address this discrepancy, the Comparative Brain Tumor Consortium has been established to study the molecular, genetic, and histologic relationship between animal and human malignancies in order to eliminate barriers in the integration of animals in all aspects of brain tumor research.^3^ The current work is thus expected to contribute to this effort.

Protein glycosylation is a ubiquitous post-translational modification found on over 50% of the proteome. Classical types of protein glycosylation are N- and O-linked, which modify Asn and Ser/Thr residues, respectively.^6,7^ N-linked glycosylation can occur when an Asn residue is found in the consensus sequon Asn_X_Ser/Thr, where X symbolizes any amino acid except Pro. N-linked glycans have a core structure consisting of 2 β-N-acetylglucosamine residues (GlcNAc2) connected to 3 mannose units (GlcNAc2-Man3), which can be extended into complex, hybrid, or high-mannose structures. These glycan structures are important in protein folding, stability, and cell-to-cell interactions outside of the cell.^8^ Extracellular O-linked glycosylation is often initiated by an α-N-acetylgalactosamine (O-GalNAc) on Ser/Thr residues and is commonly referred to as “mucin-type O-glycosylation” since it is frequently found on densely glycosylated mucin domains. O-GalNAc can be extended in monosaccharide units to create large, branched structures that are important for cell adhesion, cell polarization, and again, cell-to-cell interactions in the extracellular matrix.^8,9^ Other types of O-glycosylation include O-linked β-N-acetylglucosamine (O-GlcNAc),^10,11^ O-mannose,^12–16^ O-fucose,^17^ and O-xylose,^18^ among others.^19^ While we are only beginning to understand the latter forms of glycosylation, O-GlcNAc has been extensively described as an intracellular signaling molecule involved in cross-talk with phosphorylation.^20,21^ Finally, less commonly, β-N-acetylglucosamine has been reported to modify Cys, creating an S-linked GlcNAc structure.^22^

Aberrant glycosylation is a classic hallmark of malignant transformation; this observation has been well-documented and extensively reviewed.^8,9,23–26^ The mechanisms that produce these abnormal glycosylation patterns are broad because glycosylation is a non-template driven process and is instead reliant on expression of various glycosyltransferases, substrate availability,^27^ chaperone function,^28^ and the cellular milieu.^25^ As such, diverse glycan structures result from these dysregulated systems. For instance, altered branching,^29^ increased fucosylation,^30^ and upregulated sialylation^31^ of N-glycans has been linked to several cancerous processes,^32^ including the progression of oligodendroglioma to glioblastoma.^33^ Additionally, mucin-type O-glycans are often truncated and have been typically associated with increased cell migration and tumor metastasis, due to a loss of cell polarization.^8,9^ These shortened O-glycans comprise oncofetal antigens, neoantigens, and altered levels of normal antigens^34^ Finally, one of the most widely observed changes in glycan structures is an upregulation of sialic acid,^26,35^ which limits complement activation,^36^ engages inhibitory sialic acid-binding immunoglobulin type lectins (Siglecs),^37,38^ and reduces attachment of tumor cells to the basement membrane.^39^ These changes ultimately allow tumor cells to evade the immune system and increase invasion, causing metastasis. Most of the changes listed here are tumor-specific or tumor-associated, making them viable biomarker targets for glycan- or glycoprotein-based antibody or antibody-conjugated enzyme therapy.^40–42^

The importance of glycosylation in cancer and other pathologies has prompted the in-depth examination of the glycome and the glycoconjugates that they modify. Mass spectrometry (MS) is the premier technique to probe the proteome, glycome, and glycoproteome.^7,32^ Here we refer to *proteomics* as analysis of non- or de-glycosylated peptides/proteins, *glycomics* as the study of released or free glycans, and *glycoproteomics* as investigation of intact glycopeptides to ascertain the peptide sequence, site of modification, and glycan composition. The most common MS techniques use reverse-phase (RP)-HPLC to separate analytes followed by electrospray ionization (ESI) and mass analysis. Generally, glycopeptides are then subjected to different types of fragmentation (i.e. tandem MS) in order to glean information about the peptide and attached glycan.^7^ Some of the most common fragmentation techniques include higher energy collision dissociation (HCD),^43^ electron transfer dissociation (ETD),^44^ and supplemental activation with ETD (EThcD,^45^ AI-ETD^46^). With these techniques, it is possible to sequence a peptide, ascertain information about the glycan, and (in some cases) localize the glycosylation site.^47,48^

Another MS modality, matrix-assisted laser desorption ionization (MALDI) MS imaging (MSI), allows for the determination of the spatial distribution of glycans after enzymatic release from their protein conjugates.^49,50^ In this technique, slices from fresh-frozen or formalin-fixed, paraffin-embedded (FFPE) tissues are applied to slides, followed by antigen retrieval, the addition of de-N-glycanase (PNGaseF) to remove N-linked glycans, and coating of the tissue with matrix (e.g. 1,5-dihydroxybenzoic acid (DHB) or alpha-cyano-4-hydroxycinnamic acid (CHCA)). The images are generated by rastering a laser across the tissue slices, and the ionized glycans are generally analyzed in a time-of-flight (TOF) or ion cyclotron (ICR) mass analyzer.^49,50^ One downside of this technique is that only N-glycans can be imaged, because there is no enzyme that can perform universal release of O-glycans. Despite this drawback, the N-glycan analysis provides an estimation of the spatial distribution of the N-glycome and its changes in a tissue, disease, and/or cell type-specific manner. MALDI-MSI of N-glycans has been used on ovarian,^51^ pancreatic,^52^ and hepatocellular^53^ cancer tissue samples, and has demonstrated marked N-glycan changes in benign versus tumor regions. Further, it has also been used for the generation of a classification model for colon carcinoma tissue microarrays.^52^ Some groups have also achieved *in situ* derivatization of different sialic acid linages, ^54^ sequential PNGaseF/trypsin digestion of the same tissue,^55,56^ and visualization of extracellular matrix proteins.^57^ Taken together, MALDI-MSI is a powerful technique to image the N-glycome.

Recent developments in microextraction strategies in our laboratory coupled with MALDI-MSI have allowed us to perform spatially-resolved proteomics on various biological samples from fresh frozen and FFPE tissues.^58,59^ This spatially-resolved proteomics strategy is advantageous in that protein digestion and extraction are performed only on a restricted region of interest defined by a prior MALDI imaging experiment on a consecutive section. The strategy is essentially a solid-liquid extraction method following micro-digestion which is done by depositing picoliter quantities of enzyme using a microspotter.^60^ This directed digestion and extraction is, in essence, a means of concentrating the analyte of interest by performing digestion only in regions where this analyte is present, thereby minimizing the amount of abundant proteins extracted. The maximum spatial resolution that this strategy can attain is highly dependent on the size of the microdigested spot.^61^

In the present work, we combine the power of MALDI-MSI N-glycan imaging with our microscale proteomics technology to better understand dysregulated glycosylation in canine glioma samples. We first used standard N-glycan imaging techniques to image N-linked glycans from the surface of various canine brain tumors, finding that sialylated glycan structures were more common in the tumor and necrotic regions. We confirmed this result using Sambucus nigra (SNA) lectin staining, which selectively stains sialylated glycans. Then, we applied our microscale proteomics strategy to extract glycopeptides from tissue, which we hypothesized could be used to assign peptide and protein conjugates to better understand our N-glycan imaging results. Using this technique, we identified over 400 unique glycopeptides from 30 glycoproteins, including complex and oligomannose N-glycosylated peptides. Several of the N-glycans were also found in the MALDI MSI experiments, thus providing us with the first instance of directly linked N-glycan imaging with intact glycopeptide sequencing. Surprisingly, we also identified other glycoconjugates, including peptides modified by mucin-type O-GalNAc, O-linked GlcNAc, S-linked GlcNAc, and O-mannose. We used this information to demonstrate that there is a significant increase in sialylated O-GalNAc structures in tumor/necrotic regions compared to benign, whereas there is significantly less S- and O-GlcNAc peptides in the cancerous regions. Taken together, this proof-of-principle experiment has demonstrated that we can perform both MALDI-MSI and glycopeptide analysis on FFPE tumor tissue to better understand the glycoproteomic changes in malignant transformation.

## EXPERIMENTAL SECTION

### Reagents

HPLC grade ACN, methanol (MeOH), absolute ethanol, xylene, hydrochloric acid (HCl) and water (H2O), and analytical reagent (AR) grade TFA and sodium bicarbonate (NaHCO3) were obtained from Biosolve B. V. (Dieuze, France). 2,5 dihydroxybenzoic acid (DHB), formaldehyde, 4’,6-diamidino-2-phenylindole (DAPI), poly-D-lysine hydrobromide and bovine serum albumin (BSA) were obtained from Sigma-Aldrich (Saint-Quentin Fallavier, France). Biotech grade Tris was obtained from Interchim (Montlucon, France). Fluorescence mounting medium was obtained from Dako (Santa Clara, CA). Fluoresceine-tagged Sambuccus nigra agglutinin (SNA) was obtained as a kind gift from Prof. Anne Harduin-Lepers (Structural and Functional Glycobiology Unit, University of Lille). Glycerol-free N-glycosidase F (PNgase F, 15,000 units) was purchased from New England Biolabs (Evry, France), while sequencing-grade, modified trypsin was obtained from Promega (Charbonnières-les-Bains, France).

### Biopsies and tissue preparation

The study was performed with the approval and in accordance to the guidelines of the Oncovet ethical committee. Biopsies were taken from dog patient samples at the Oncovet Clinic with the approved consent of their owners. The biopsies were immediately subjected to formalin-fixed, paraffin-embedding using standard protocols. An overview of the characteristics of dog patients and collected biopsies can be found in **Supplementary Data 1**.

### MALDI MSI

Sections (8 μm thick) were taken from the FFPE blocks using a microtome (Leica Biosystems, Nanterre, France) and mounted on indium/tin oxide-coated slides (ITO, LaserBioLabs, Sophia-Antipolis, France). The ITO slides were pre-treated by pipetting 1.5 mL of polylysine solution on the conductive surface and incubating at room temperature for 5 min. The treatment was performed twice and dried under a heat gun then dipped in HPLC water. The mounted sections were heated at 60 °C on a slide heater for 1 h. While the slides were still hot, the sections were dewaxed in xylene and rehydrated following standard procedures.^57^ The slides were then subjected to antigen retrieval by incubating in 20 mM Tris-HCl (pH = 9) at 95 °C for 1h, followed by rinsing in HPLC water. PNGaseF (New England Biosystems) was subjected to dialysis by pipetting 40 μL of PNGaseF onto a PVDF membrane at set on top of 200 mL of water. The buffer exchange was allowed to proceed for 2 h at RT. The PNGaseF was then rehydrated and sprayed onto tissue slices at a rate of 10 μL/min for a total of 15 layers. The slides were placed in a humidified chamber and left overnight at 37 °C. After the reaction, DHB matrix was sublimated onto the sections at 140 °C for 10 min, which leads to an average of 0.424 mg matrix deposited per cm^2^ based on three independent measurements.

MS imaging of tissue slices was performed using a RapiFlex MALDI TOF instrument (Bruker Daltonics, Bremen, Germany). Images were acquired at 70 μm resolution scanning at m/z 700-3,200 with the Smartbeam 3D laser firing at a frequency of 10 kHz. Each spectrum was recorded after accumulating 1,000 laser shots per spot. The laser ablation pattern was set at M5 at a 35 x 35 μm scan range. The ion source voltage was set to 19.984 kV, while the PIE and lens were at 2.608 and 12.366 kV, respectively. The reflectors were set at 20.663, 1.112 and 8.558 kV. Matrix suppression by deflection up to m/z 240 was activated. The reflector detector gain and sampling rate were kept constant on all imaging acquisitions.

The Glycomod tool (https://web.expasy.org/glycomod/) was initially used to predict the structure and composition of the observed N-glycans from the full MS spectra recorded at high mass accuracy, with the searches performed at 0.1 Da mass tolerance. Potential glycan hits were matched with the masses detected in the MALDI MS images. The MS images were uploaded on SCiLS Lab version 2016b (Bruker Daltonics) and the ion distributions (as sodiated adducts or oxidized forms) of the glycans were mapped by plotting the signal intensities in all spectra normalized against the total ion count. To confirm the glycan structure predicted by GlycoMod, PNGaseF-released samples were subjected to MSn analysis using a MALDI LTQ orbitrap XL instrument (ThermoFisher Scientific, Bremen, Germany) operated at 30,000 FWHM at m/z 400. Glycans were detected as sodiated adducts in positive mode. Spectra were acquired directly on-tissue and were averaged from 10 scans with each scan composed of 1 μscan acquired at 1 μscan/step and 10 laser shots.

### MSn

MSn spectra were acquired directly on tissue using a MALDI LTQ Orbitrap instrument after images had been obtained. The MALDI LTQ Orbitrap XL is equipped with a commercial N2 laser (LTB Lasertechnik, Berlin, Germany) operating at λ = 337 nm with a maximum repetition rate of 60 Hz. The hybrid configuration replaces the heated capillary of the electrospray source with a q00 that sends packets of ions into a linear trap for collision-induced fragmentation (CID), with the fragment ions then being concentrated in a C-trap and transferred to the orbitrap for high-resolution mass analysis. The maximum energy per pulse was set to 12 μJ. Precursor ion isolation was performed using an isolation window between ±1 and ±3 Da and the fragments scanned with a maximum accumulation time of 120 ms. Succeeding MSn of the daughter ions were performed with a maximum accumulation time of 180 ms. External calibration was performed using the ProteoMass MALDI Calibration Kit (Sigma-Aldrich, St. Quentin-Fallavier, France).

### HES and Lectin staining

HES staining was performed as previously described and slices were scored by a licensed veterinary pathologist (**Supplementary Data 1** for more information). For lectin staining, using a Dako delimiting pen (Agilent Technologies, Santa Clara, CA), dams were created around tissue sections. The sections were then incubated in 1% BSA (w/v) in approximately 300 μL of PBS for 30 min at RT, then incubated in 10 μg/mL of SNA lectin for 2 h. They were then rinsed for 10 min three times with 1% BSA in PBS. The sections were then incubated in approximately 300 μL of DAPI for 20 min and rinsed with PBS for 5 min. Finally, two drops of Vectashield fluorescence mounting medium (Dako, Agilent Technologies) was added and the sections were cover-slipped and sealed with nail polish. Confocal images were obtained using a fluorescence microscope (Leica Biosystems). Adjacent tissue sections incubated in 1% BSA in PBS served as controls. Zeiss LSM700 confocal microscope connected to a Zeiss Axiovert 200 M with an EC Plan-Neofluar 40x/1.30 numerical aperture oil immersion objective (Carl Zeiss AG, Oberkochen, Germany). Processing of the images was performed using Zen software and applied on the entire images as well as on controls. The presented pictures are representative of independent triplicates.

### On-tissue microdigestion

Regions of interest (ROIs) discerned using MALDI-MSI, HPS staining, and SNA lectin staining were marked on adjacent tissue sections and optical scans were obtained for reference. Five spots were marked per ROI, and these were digested by microspotting 20 μg/mL of trypsin suspended in 50 mM NH4HCO3 using a chemical inkjet printer (CHIP 1000, Shimadzu, Kyoto, Japan). Each spot was composed of 4 microspots spaced 100 μm center-to-center, with each microspot produced by printing 13 droplets at 200 pL/droplet. By optimizing the waiting time between each pass, this yielded spots with diameters between 450-600 μm. The spots were maintained wet for 2 h, after which, the sections were incubated at 37 °C inside an enclosed glass chamber humidified with 1:1 MeOH/H2O for 1h. The sections were then dried under vacuum for 5 min.

The following solutions were used to extract the digested peptides: 0.1% TFA in water, 4:1 ACN/0.1% TFA in water, and 7:3 MeOH/0.1% TFA in water. Each solution (3 μL) was deposited to cover all the digested spots for one region, and the extracts were manually pipetted 10x before recovery. This was repeated once before proceeding with the next solvent system. In cases where the spots were distant from each other, the total volume was divided per spot and extraction was performed separately. The extracts were then frozen in −80 °C and dried using a speedvac. The dried extracts were reconstituted in 10 μL 0.1% TFA in water, vortexed for 10 s and sonicated for 5 min. They were then desalted using C18 Ziptips (Pierce, Thermo Fisher Scientific) and taken to dryness in a vacuum concentrator.

### Glycoproteomic MS analysis

All of the glycoproteomic samples were analyzed by LC-MS/MS on an Orbitrap Fusion Tribrid (Thermo Fisher Scientific) coupled to a Dionex Ultimate 3000 HPLC. The samples were reconstituted in 7 μL of 0.1% formic acid in water (“buffer A”). Then, a portion of the sample (6.5 μL) was loaded via autosampler isocratically onto a C18 nano pre-column using 0.1% formic acid in water (“Solvent A”). For pre-concentration and desalting, the column was washed with 2% ACN and 0.1% formic acid in water (“loading pump solvent”). Subsequently, the C18 nano pre-column was switched in line with the C18 nano separation column and injected at 0.3 μL/min onto a 75 μm x 250 mm EASY-Spray column (Thermo Fisher Scientific) containing 2 μm C18 beads. The column was held at 40 °C using a column heater in the EASY-Spray ionization source (Thermo Fisher Scientific). The samples were eluted at 0.3 μL/min using a 90-min gradient and a 185-min instrument method. Solvent A was comprised of 0.1% formic acid in water, whereas Solvent B was 0.1% formic acid in acetonitrile. The gradient profile was as follows (min:%B) 0:3, 3:3, 93:35, 103:42, 104:98, 109:98, 110:3, 185:3. The instrument method used an MS1 resolution of 60,000 at FWHM 400 m/z, an AGC target of 3e5, and a mass range from 300 to 1,500 m/z. Dynamic exclusion was enabled with a repeat count of 3, repeat duration of 10 s, exclusion duration of 10 s. Only charge states 2-6 were selected for fragmentation. MS2s were generated at top speed for 3 s. HCD was performed on all selected precursor masses with the following parameters: isolation window of 2 m/z, 30% collision energy, Orbitrap detection with a resolution of 30,000, and an AGC target of 1e4 ions.

### Glycoproteomic data analysis

Raw files were searched using Byonic by ProteinMetrics against the Uniprot *Canis familiaris* database (downloaded November 2018). Search parameters included semi-specific cleavage specificity at the C-terminal site of R and K. Mass tolerance was set at 10 ppm for MS1s, 0.1 for MS2s. Methionine oxidation (common 2), asparagine deamidation (common 2), and N-term acetylation (rare 1) were set as variable modifications with a total common max of 3, rare max of 1. Glycosylation was added in three separate searches to minimize search times. In the first search, N-glycans were set as variable modifications (common 2), using the “N-glycan 57 human plasma” database. In the second iteration, O-glycans were set as variable modifications (common 2), using the “O-glycan 6 most common” database. In the final search, an O-mannose database containing (Hex, Hex-HexNAc, Hex-HexNAc2, Hex2-HexNAc-NeuAc, and Hex2-HexNAc-Fuc) was used for a variable modification (common 2). Cysteine carbaminomethylation was set as a fixed modification. Peptide hits were filtered using a 1% FDR. All peptides were manually validated and/or sequenced using Xcalibur software (Thermo Fisher Scientific). For statistical analyses, one-way ANOVAs were performed for various glycan counts comparing benign, tumor, and necrotic conditions. For instances where p<0.05, Tukey’s Multiple Comparisons post-hocs determined significantly different comparisons.

### Subnetwork enrichment pathway analyses and statistical testing

The raw files described above were also searched using MaxQuant with the following parameters: fixed cysteine carbaminomethylation, variable deamidation of asparagine, and variable methionine oxidation. The Elsevier’s Pathway Studio version 11.0 (Ariadne Genomics/Elsevier) was used to analyze relationships among differentially expressed proteomics protein candidates using the Ariadne ResNet database.^62,63^ “Subnetwork Enrichment Analysis” (SNEA) algorithm was selected to extract statistically significant altered biological and functional pathways pertaining to each identified set of protein hits among cluster 1 (overexpressed in tumor and necrotic regions) and cluster 2 (overexpressed in benign). SNEA utilizes Fisher’s statistical test to determine if there are nonrandom associations between two categorical variables organized by specific relationship.^64^ Integrated Venn diagram analysis was performed using “the InteractiVenn”: a web-based tool for the analysis of complex data sets.^65^ See **Supplementary Data 2** for the listed differentially expressed pathways. Each table indicates the Entity designation, Relationship type, and Reference type.

## RESULTS

### MALDI-MSI shows sialylated glycans are enriched in necrotic regions

**Figure 1A** shows the ion distribution of the summed intensities of sodiated sialic acid-containing N-glycans HexNAc4-Hex5-NeuAc2, HexNAc4-Hex5-NeuAc-NeuGc and HexNAc4-Hex5-NeuGc2 and their di- and tri-sodiated adducts across different glioma biopsies. The ion distribution of the individual glycoforms is shown in **Supplementary Data 3**. Superposition of the summed ion distributions with the HES-stained section shows that on both anaplastic oligodendroglioma and glioblastoma samples, these glycoforms are present in both tumor and necrotic regions but are highly abundant in the latter (**Figure 1B**). In the anaplastic oligodendroglioma section (**Figure 1B**, left panel), the distribution extends through the pseudo-glomerular vessels present along the margins of the tumor (indicated by dark blue arrows), suggesting its possible relationship with vigorous and abnormal angiogenesis associated with high-grade glioma. However, regions where both tumor and necrosis are present only show weak distribution (indicated by green arrows). The same observation applies to the glioblastoma section shown in **Figure 1C**, right panel. In this case, however, the glycoforms are also detected in the pseudo-palisading (indicated by yellow arrows). Additionally, a plot of the normalized total ion current (TIC) intensity from spectra taken in the necrotic regions of all tissues, defined based on the HES images, confirms this finding (**Figure 1C**).

**Figure 1.**
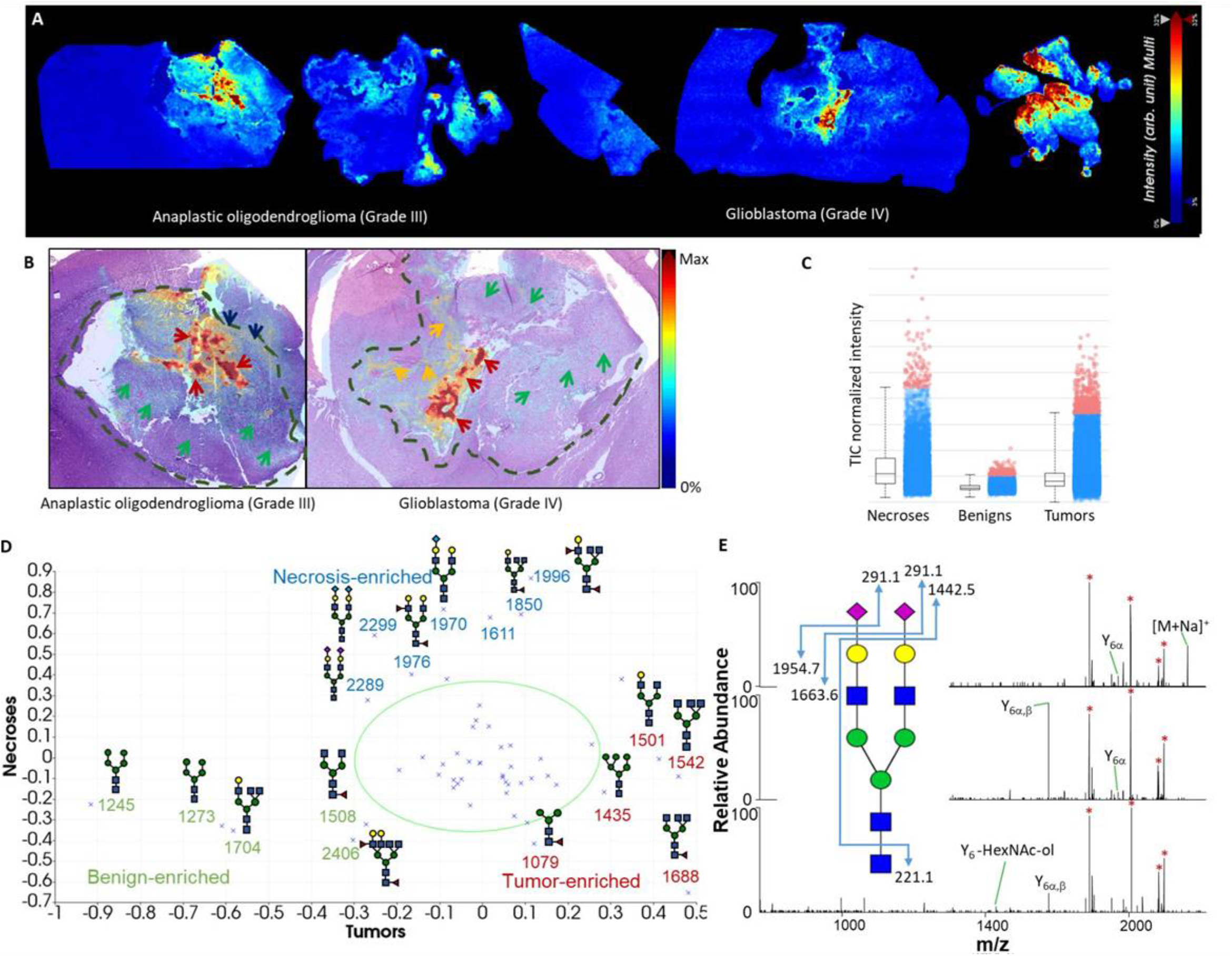
MALDI-MSI of N-glycans. A) Summed ion images of sodiated HexNAc4-Hex5-NeuAc2, HexNAc4-Hex5-NeuAc-NeuGc and HexNAc4-Hex5-NeuGc2 and their di- and tri-sodiated adducts on canine glioma biopsies. Images were generated using SCiLS Lab. B) Superposition of MALDI-MSI glycan images with H&E-stained adjacent sections. Arrows and dashed lines indicate regions annotated by the pathologist (red, green, yellow and dark blue arrows for necrotic and tumor regions, pseudo-glomerular vessels and pseudopallisading necroses, respectively, and dashed lines for tumor margins). C) Normalized intensity of total ion signals of the combined mono-, di- and tri-sodiated species from necrosis, benign, and tumor samples. D) PCA analysis of detected glycans in MSI analysis, with benign, necrosis, and tumor enriched glycans highlighted in green, blue, and red, respectively. E) MSn spectra of HexNAc4-Hex5-NeuAc2 confirming its structure.

The acquired images were exported in SCiLS and principal component-linear discriminant analysis (PC-LDA) was performed. This supervised analysis was done by pre-defining the ROIs where spectra will be taken using histological annotations from the HES optical images. The loadings plot results (**Figure 1D**) identified m/z intervals attributed to the aforementioned sialic acid-containing glycoforms (HexNAc4-Hex5-NeuAc2, m/z 2289 corresponding to [M-H+3Na]^+^ and HexNAc4-Hex5-NeuGc2, m/z 2299 corresponding to [M-H+2Na]^+^), as well as other complex, sialic acid-containing glycoforms HexNAc3-Hex3-Fuc-NeuAc (m/z 1611, structure unassigned) and HexNAc4-Hex5-NeuAc (m/z 1970). The loadings plot also identified m/z enriched in tumor samples, including the fucosylated complex glycoforms HexNAc5-Hex3-Fuc (m/z 1688) and HexNAc5-Hex3-Fuc (m/z 1542). On the other hand, notable glycoforms enriched in the benign regions include the high mannose-type HexNac2-Hex5 (m/z 1257 and its oxidized form m/z 1273) as well as fucosylated complex glycoforms HexNAc4-Hex3-Fuc (m/z 1508) and HexNAc6-Hex5-Fuc2 (m/z 2406).

To confirm the glycoform assignment based on GlycoMod searches, we repeated the mass measurements on an LTQ-Orbitrap XL instrument equipped with a MALDI source (**Supplementary Data 4**). Collision-based MSn fragmentation (**Figure 1B**) was performed where possible. MS2 of the precursor ion of HexNAc4-Hex5-NeuAc2 ([M+Na]^+^, m/z 2245) leads to sequential losses of the glycan residues, starting with the first sialic acid producing the Y6α ion with mass similar to HexNAc4-Hex5-NeuAc (m/z 1954.596, **Figure 1B**, top panel). This is followed by the loss of the second sialic acid, yielding the Y6α,β ion with a mass similar to HexNAc4-Hex5 (m/z 1663.580, middle panel). MS4 then leads to loss of the terminal GlcNAc (m/z 221.088) yielding the Y6-HexNAc-ol ion. Structures of other glycoforms were also partially elucidated using their MSn spectra and are shown in **Supplementary Data 2**, thus confirming their glycan identities as predicted by GlycoMod. The list of all glycoforms detected in the glioma samples is provided in **Supplementary Data 5**.

### SNA lectin staining reveals sialic acid-containing glycans in other tissue regions

In order to further verify the distribution of sialylated glycan structures in the necrotic region(s), SNA lectin staining of consecutive slices was performed. Results for all of the biopsies are shown in **Supplementary Data 6**. Confocal images taken of the different regions of WHO grade III (**Figure 2A**) and grade IV (**Figure 2B**) biopsies, demonstrated that sialic acid was present in necrotic zones. Tumor regions and regions marked by the presence of both tumor and necrosis show positive staining, concomitant with findings from the MALDI imaging experiments. One glioblastoma sample (**Figure 2B**, bottom panels) showed very intense staining in the tumor region, although the MS image only shows moderate distribution of sialic acid-containing glycans in this region. Benign regions particularly the corpus callosum (**Figure 2A**, bottom panels) and choroid plexi (**Supplementary Data 6**, last slide) likewise show intense staining, although no sialic acid-containing N-glycans were detected in these zones in MALDI imaging experiments. SNA stains for 2,6-linked sialic acids and does not discriminate with regard to their origin. Thus, we suspected that the abundant SNA signal was derived from other sources, such as O-linked glycans. To investigate this hypothesis, we turned to the microscale proteomics technique our laboratory developed.

**Figure 2.**
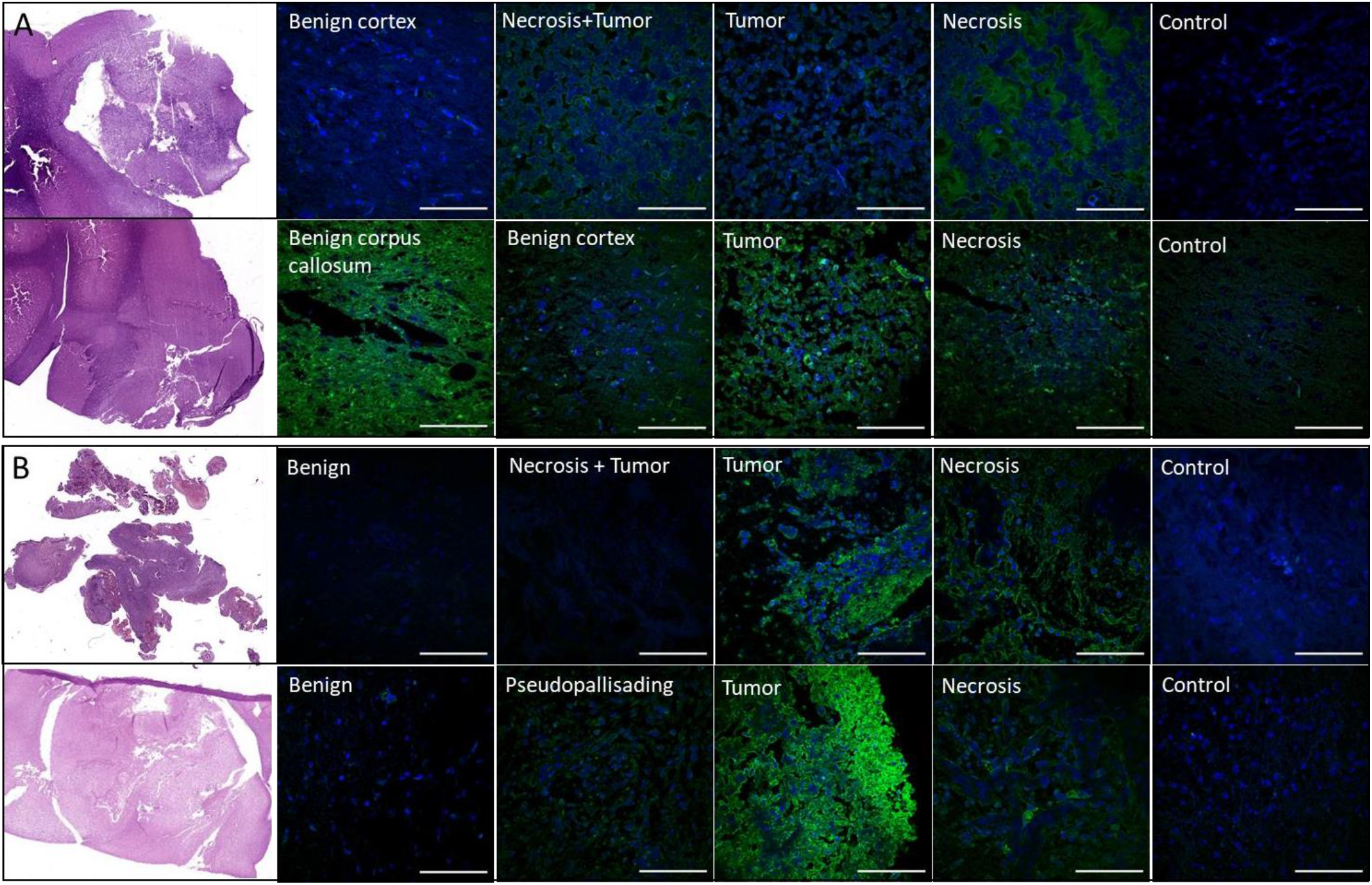
SNA staining of tissue slices. Confocal fluorescent images taken at pathologist-annotated regions present in oligodendroglioma (WHO grade III, A) and glioblastoma (WHO grade IV, B) samples. HES stains (left) present for comparison to fluorescent images (right). The sections were incubated in 1% BSA (w/v) in 300 μL of PBS for 30 min at RT, then incubated in 10 μg/mL of SNA lectin for 2 h. Slides were then rinsed for 10 min 3x with 1% BSA in PBS, then incubated in approximately 300 μL of DAPI for 20 min. Confocal images were obtained using a fluorescence microscope (Leica Biosystems). Images taken from separate adjacent sections that were incubated without the lectin serve as controls.

### Spatially-resolved proteomic analysis reveals hierarchical clustering of benign and cancerous tissue

Regions of interest (ROIs) identified using both N-glycan MALDI imaging and SNA lectin staining were subjected to microdigestion by depositing picoliters of trypsin using a microspotter (**Figure 3A**). We began with 8 biopsy tissues which resulted in the microdigestion of 11 benign, 4 margin, 11 tumor, and 7 necrosis samples. The samples were subjected to LC-MS/MS on a Thermo Orbitrap Fusion instrument with higher-energy collision dissociation (HCD), and the raw files were searched using MaxQuant (unmodified peptides) and Byonic (glycopeptides). Hierarchical clustering of the protein groups identified by MaxQuant using the Andromeda search engine and quantified using LFQ intensities shows that samples from the benign regions cluster distinctly from those obtained from the tumor and necrotic zones (**Figure 3B**), whereas samples taken from the margins do not form a distinct cluster. The overexpressed proteins of the “benign” cluster are associated with pathways related to synaptic processes such as synaptogenesis, synaptic plasticity, synaptic vesicle endo- and exocytosis, transport, docking and fusion, and long-term synaptic potentiation and depression (**Figure 3C**). On the other hand, overexpressed proteins in the “tumor and necrosis” cluster are associated with cancer processes such as cell proliferation, growth, division, metastasis and migration, as well as RNA splicing, apoptosis and neo-plastic growth (**Figure 3D**).

**Figure 3.**
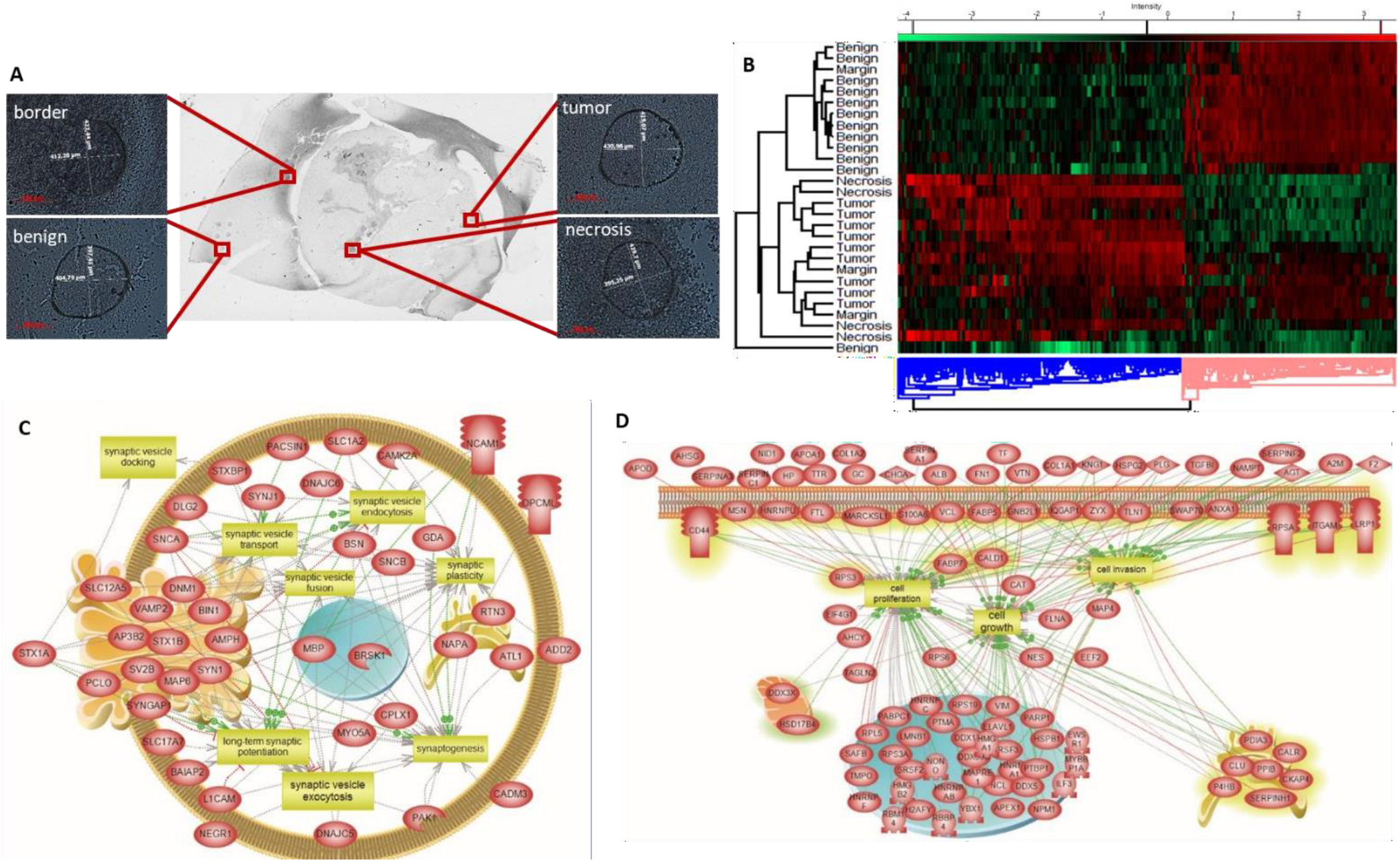
Spatially-resolved proteomic analysis of tissue slices. A) Zoomed optical images of sample microspots from each ROI after microdigestion using the CHIP-1000 printer. B) Hierarchical clustering of protein identifications with ANOVA-significant (p < 0.01) differential expression across the different ROIs, using the LFQ intensities calculated by MaxQuant. C) Selected overrepresented pathways in the “benign” cluster (pink, Figure 3B). D) Selected overrepresented pathways in the “necrosis and tumor” cluster (blue, Figure 3B).

### Spatially-resolved glycoproteomic analysis links MSI glycan imaging with intact N-glycopeptide identification

The microdigested areas were then extracted with different solvent mixtures to maximize glycopeptide extraction. Glycoproteomic analysis identified the glycopeptide MVSHHnLTSGATLINEQWLLTTAK from haptoglobin bearing the HexNAc4-Hex5-NeuAc2 motif (**Figure 4A, Supplementary Data 7 and 8**.). Non-glycosylated peptides of haptoglobin were also detected, both in necrotic as well as other regions, but the glycosylated peptide was only observed in necrotic regions. Unfortunately, this was the only sialylated N-glycopeptide that we observed. Other N-glycopeptides detected include those bearing high mannose motifs from extracellular matrix (ECM) chondroitin sulfate proteoglycans such as neurocan (AnATLLLGPLR, **Figure 4B**), members of the immunoglobulin superfamily such as neurofascin (IgCAMs), limbic system-associated membrane protein (LAMP, IgLONs), contactin 1 (contactin CAMs), Thy-1 cell surface antigen (CD90), and neuroprotective factor prosaposin. Interestingly, while overall numbers of N-glycopeptides were similar between various regions, the HexNAc2-Hex5 glycan was significantly enriched (66 glycopeptides) in the benign regions when compared to tumor (18 glycopeptides, p<0.01**) and necrotic regions (25 glycopeptides, p<0.05* for necrosis; **Figure 5A and B**). This correlates with our MALDI MSI data, which demonstrated a marked increase in HexNAc2-Hex5 glycans in benign regions. Finally, we observed several glycans modified by fucose, as demonstrated in **Figure 4C**, with a glycopeptide from the antimicrobial protein myeloperoxidase. No significant change between benign and cancerous fucosylation was observed.

**Figure 4.**
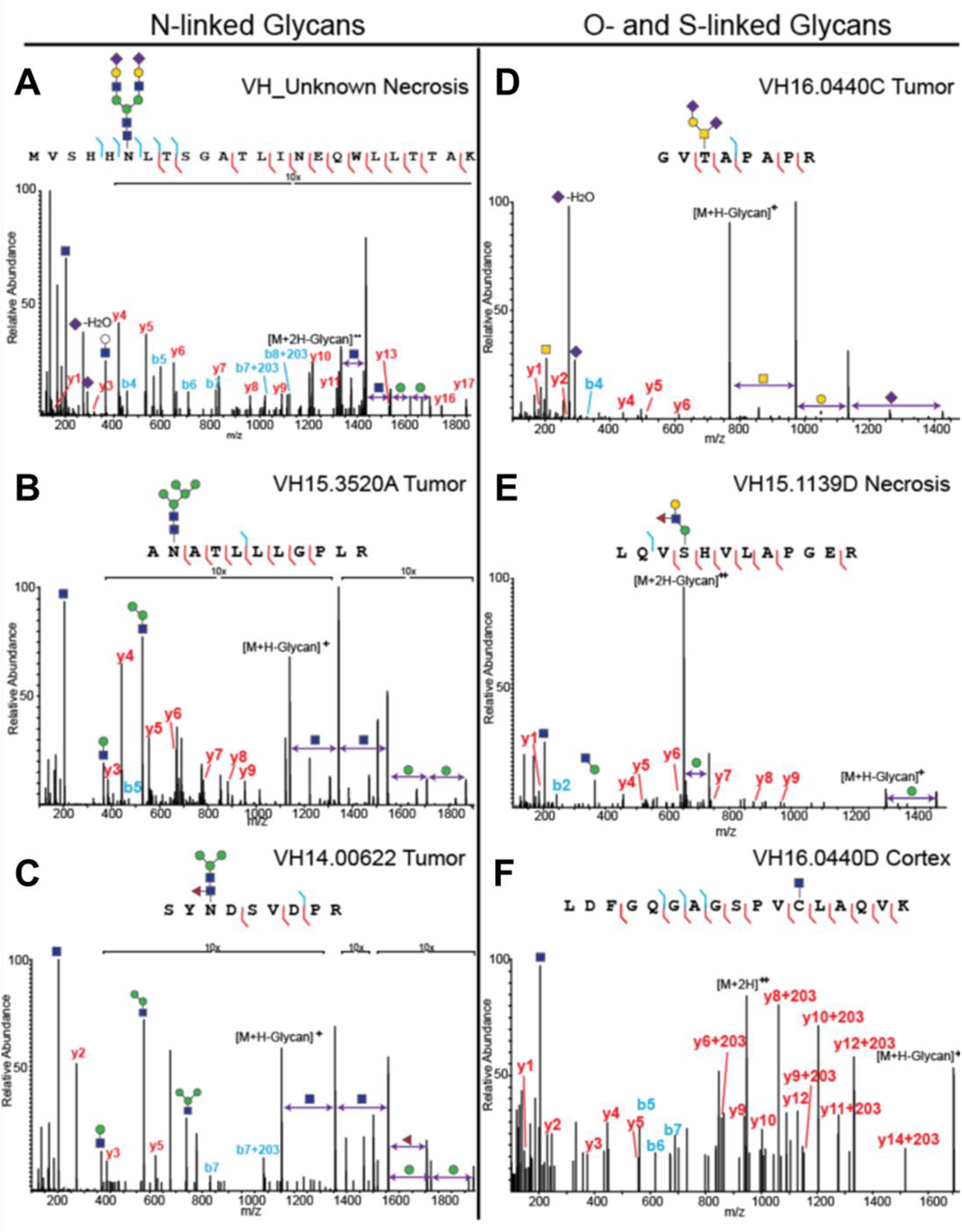
Spatially-resolved glycoproteomics identifies several types of glycosylation. All microdigested samples were subjected to LC-MS/MS analysis on a Thermo Orbitrap Fusion Tribrid, and peptides were subjected to HCD fragmentation. Note that in the case of O-glycosylation, the site of modification is only localized because there is only one possible site of modification. Spectra were annotated manually to confirm glycan composition, peptide sequence, and (if possible) site localize the glycan. A) Haptoglobin peptide MVSHHnLTSGATLINEQWLLTTAK bearing the complex, disialylated glycan HexNAc4-Hex5-NeuAc2. B) Peptide from neurocan, AnATLLLGPLR, modified with an N-linked high mannose (HexNAc2-Hex5) structure. C) Myeloperoxidase peptide SYnDSVDPR modified with a fucosylated paucimannose N-glycan (HexNAc2-Hex3-Fuc). D) Protein tyrosine phosphatase receptor Z1 peptide GVtAPAPR modified with a disialylated, core 1 structure (HexNAc-Hex-NeuAc2). E) Another protein tyrosine phosphatase receptor Z1 peptide, LQVsHVLAPEGR, modified with an extended O-mannose glycan (Hex2-HexNAc-Fuc). F) Bassoon presynaptic cytomatrix protein peptide LDFGQGAGSPVcLAQVK was modified by an S-linked GlcNAc.

**Figure 5.**
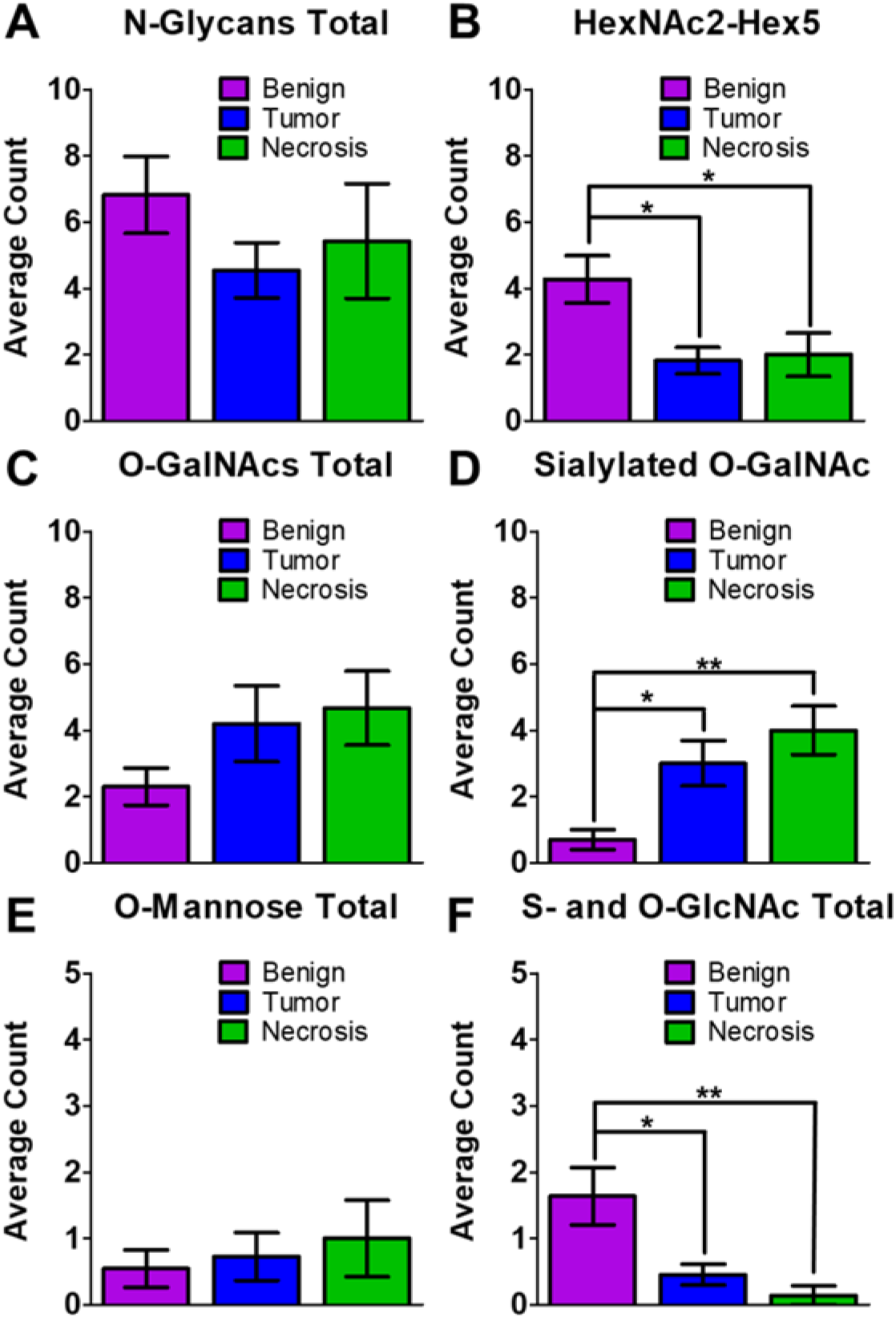
Statistical analysis of glycan changes between samples. For all samples, a 3-way ANOVA test was performed in GraphPad PRISM, wherein (*) indicates a p value of <0.05, and (**) indicates a p value of <0.01. A) The total number of N-glycopeptides was not found to be significantly different between the three types of samples. B) The number of glycopeptides modified by HexNAc2-Hex5 was found to be significantly higher in the benign samples compared to both tumor and necrotic regions. C) The total number of O-GalNAcylated peptides was not was not found to be significantly changed between the three regions. D) The number of sialylated O-GalNAc glycopeptides was significantly increased in tumor and necrotic regions when compared to benign. E) The total number of O-Mannosylated peptides was not was not found to be significantly changed between the three regions. F) S- and O-linked GlcNAcylated peptides were significantly decreased in both tumor and necrotic regions when compared to benign.

### Identification of diverse glycopeptides from spatially-resolved glycoproteomic experiments

This work was initially focused on finding glycoconjugates modified by the same N-glycans that we observed in MALDI MSI experiments. However, it was clear after SNA staining and the N-linked glycopeptide analysis that there were other glycosylated species in the samples. Thus, we re-analyzed the data for O-linked mucin-type glycans, which cannot be imaged by MALDI MSI due to the harsh chemical conditions necessary to release them. We detected a surprisingly large number of O-GalNAcylated peptides in the samples, all of which can be found in **Supplementary Data 7** and **8**. Most of the glycans that we detected were tumor associated carbohydrate antigens (TACAs), such as the T-antigen (GalNAc-Gal) and the sialylated T-antigen (GalNAc-Gal-NeuAc). One example of this is shown in **Figure 4D**; the glycopeptide GVtAPAPR from protein phosphatase receptor type Z1 (PTPRZ1, also called phosphacan) was modified by a disialyl-T antigen (GalNac-Hex-NeuAc2) and was found in 5 of 8 tumor biopsies, in 3 of 5 necrosis biopsies, and in all margin regions. The same glycopeptide was also detected with the T-antigen or sialyl-T antigen in some tissue regions (**Supplementary Data 7 and 8**). Mono- and disialylated core 1 O-glycans were also highly expressed on ECM lecticans including brevican, versican, and neurocan, and were also observed in fibronectin, fibrinogen, and apolipoprotein E. While we did not detect a statistically significant change in O-GalNAcylated peptides between the benign and tumor/necrotic sections, we did observe a significant increase in sialylated O-GalNAc peptides in the tumor (p<0.01 **) and necrotic (p <0.01 **) regions (**Figure 5C** and **D**). Finally, while many of the proteins we observed have been noted previously as O-glycoproteins, neurocan and oligodentocyte myelin glycoproteins had not. To the best of our knowledge, this is the first observation of O-glycosylation on these proteins.

Further analysis of the glycoproteomic data revealed the presence of O-mannosylated glycopeptides, which is perhaps unsurprising because O-mannose accounts for up to 30% of brain O-glycans.^13,14,16^ An example is shown in **Figure 4E**, which displays an annotated glycopeptide spectrum from PTPRZ1. The sequence LQVsHVLAPEGR bears a fucosylated core M1 type O-mannose (Hex2-GlcNAc-Fuc) that contains a Lewis X structure. This glycopeptide was observed in 3 of 11 benign samples, 2 tumor and necrosis samples, and 1 margin sample. PTPRZ1 was previously identified as a substrate for O-mannosyl glycosylation by O-mannose β-1,2-N-acetylglucosaminyltransferase (POMGnT1).^66^ We also observed a peptide from cadherin 13, another known target of O-mannosylation.^12^ The three other proteins that we found to be O-mannosylated have heretofore never been described as O-mannosylated and are thus novel modifications. These proteins include albumin, collagen alpha-1 chain, and KH-type splicing regulatory protein (**Supplementary Data 7** and **8**). We did not detect a statistically significant change in O-mannosylated peptides between benign and tumor/necrotic regions (**Figure 5E**).

Finally, intracellular O-GlcNAc glycopeptides were also observed, and were detected mainly on synapsin-1 and spectrin beta chain. Synapsin-1 was detected only in the benign regions, while spectrin peptides were observed in the benign, margin, and tumor regions. Additionally, a glycopeptide (LDFGQGAGSPVcLAQVK) bearing the rare S-linked glycan attached to a Cys residue, was detected predominantly in benign regions (7 out of 11 samples, **Figure 4E**). This glycopeptide is part of the bassoon presynaptic cytomatrix protein, a structural protein found in the ribbon synapse, and is involved with presynaptic vesicle release.^67^ The same S-GlcNAcylated glycopeptide, as well as others from bassoon, were described recently by Burlingame and co-workers in the mouse synaptosome.^22^ Interestingly, and contrary to several other reports, we observed a significant decrease in O- and S-linked GlcNAcylation in the tumor (p<0.05 *) and necrotic (p<0.01 **) regions when compared to benign tissues (**Figure 5F**). Taken together, these data demonstrate that our spatially-resolved glycoproteomics technique can (a) link MALDI-MSI glycan imaging experiments to the associated glycoconjugate (**Figure 4A-C**), (b) allow for the detection of a wide array of glycopeptide modifications (**Figure 4**), and (c) distinguish diverse and significant glycan changes in benign, tumor, and necrotic regions (**Figure 5**).

## DISCUSSION

Malignant glioma is the most common primary tumor in the central nervous system and has an extremely poor prognosis. Canine glioma is a good model system for human brain cancer because of their similar characteristics, including: spontaneous origin, high level of invasiveness, and poor clinical outcomes.^3,4^ Since glycosylation is an aberrant feature of all cancers but has been studied infrequently in glioma, this work was initially aimed at developing a MALDI-MSI workflow to visualize N-linked glycans from canine glioma FFPE tissue. We based our protocol on several published work,^49,50^ which allowed us to easily attain this goal. We noted that a biantennary complex N-linked glycan (HexNAc4-Hex5-NeuAc2) was upregulated in necrotic regions, and a high-mannose N-linked glycan (HexNAc2-Hex5) was enriched in benign regions.

Although mapping of the spatial distribution of N-glycans using MALDI MSI is useful in many aspects, the information that it can currently provide is limited. By nature of the technique, the N-glycans are removed from the proteins that they modify, thus losing the intact N-glycopeptide/protein information. In an effort to address this limitation, Heijs and co-workers developed a successive trypsin protocol after an N-glycan imaging experiment to determine which peptides have been deamidated after PNGaseF treatment.^55^ This provided insight into which proteins had been de-N-glycosylated; however, it does not directly link the detected N-glycans with their protein conjugates. This information can only be obtained if the intact glycopeptide itself has been detected and sequenced. Additionally, the technique is limited to N-glycans, since these are the only glycans that can be easily and enzymatically liberated from the protein conjugate to be visualized by MALDI-MSI.

Thus, we reasoned that our spatially-resolved proteomics method, guided by MALDI-IMS MSI glycan imaging, could potentially allow us to explain the global N-glycosylation changes observed in glioma histological specimens. In these experiments, the importance of spatial localization has to be emphasized. This is exemplified by haptoglobin, where many non-glycosylated peptides were found in all of the tissue sections we studied. However, the glycopeptide MVSHHnLTSGATLINEQWLLTTAK (N107 modified by HexNAc4-Hex5-NeuAc2) from this protein was only detected in the necrotic region, which is also where the glycan HexNAc4-Hex5-NeuAc2 was found to be upregulated in MALDI-MSI experiments. Taken together, we have demonstrated the first experiment that directly links glycan imaging with intact glycopeptide identification. Importantly, haptoglobin is a homodimer involved in the scavenging of iron Fe (III) and haptoglobin glycosylation is becoming a major target in cancer research.^68^ Canine haptoglobin contains 3 putative N-glycosylation sites and shares conserved sequences with other mammals, including humans. Specifically, N107 of canine haptoglobin shares identical flanking amino acids to N184 of human haptoglobin, which has also been shown to be modified by biantennary complex N-glycans.^68,69^

Conflicting results of SNA lectin staining with N-glycan MS imaging suggested that other types of glycans could be present on the tissues, since SNA lectin does not discriminate between 2,6-NeuAcs derived from N- and O-glycans or glycolipids. In addition to performing Byonic N-glycopeptide searches, the raw files were exhaustively searched for other potential glycopeptides that were missed by the initial search, and these were manually annotated. Results revealed a diverse set of glycopeptides including intracellular O-GlcNAc, extracellular O-linked mucin-type glycans, O-mannose, and, surprisingly, even the rare S-GlcNAc. The diversity of the structures detected highlights the importance of using complementary tools to examine canine glioma in addition to N-glycan imaging, and also emphasizes the need for development of other glyco-hydrolases for this approach.

More importantly, the on-tissue spatially-resolved glycoproteomics results provided insight into dysregulated glycosylation in glioma. For instance, brevican is the most abundant lectican in the central nervous system and is known to play a role in glioma invasiveness and cell motility.^70,71^ In particular, it has been shown that brevican has two glioma-specific isoforms - B/b_sia_ and B/b_Δg_ - that are generated by differential glycosylation and are absent from normal adult brain. While B/b_Δg_ is an underglycosylated proteoform, B/bsia is an oversialylated proteoform expressed by half of the high- and low-grade gliomas that the authors analyzed.^72^ Interestingly, though, the authors did not investigate which sites may be modified by the sialylated glycans, leaving it ambiguous as to whether the glycosylation was N- or O-linked. In this study, we report four peptides from brevican modified by O-GalNAc glycans, several of which are modified by sialic acid (**Supplemental Data 7 and 8**). We did not find any brevican peptides modified by any other type of glycosylation. Thus, our data suggests that the B/bsia proteoform may result from an increase in sialylation of O-GalNAc mucin-type glycosylation.

We also detected several other members of the hyaluronan-binding chondroitin sulfate proteoglycan family (of which brevican is a member), including phosphacan (a splice variant of PTPRZ1), neurocan, and versican. In particular, neurocan and phosphacan bind to neurons and are potent inhibitors of neuronal and glial adhesion. As such, it has been established that the upregulation of both neurocan and phosphacan have negative prognostic implications in glioblastoma.^73^ However, the in-depth impact of their glycosylation patterns remains to be elucidated, especially in the context of glioma. To the best of our knowledge, neurocan was previously unknown to be modified by mucin-type O-glycans. Here we show that neurocan is modified on at least 3 different sites by O-GalNAc glycans, and that the majority of the glycans found in tumor and necrotic regions bear sialic acid. As increased sialic acid is linked to increased invasion and metastatic potential, in part due to electrostatic repulsion, it follows that increased sialylation of these proteoglycans could be a mechanism for the characteristically high invasion of glioma.^25^

On the other hand, O-GalNAcylation and O-mannosylation of phosphacan has been reported using the SimpleCell technology developed by Clausen and colleagues.^15,74^ We corroborate their results in a glioma system, demonstrating that phosphacan is modified in several locations by various O-GalNAc and O-mannose glycans. Defects in O-mannosylation lead to abnormal neuronal migration, and have been associated with a range of muscular dystrophies collectively called α-dystroglycanopathy.^16^ α-dystroglycan also forms complexes with ECM glycoproteins particularly laminin, and aberrant O-mannosylation of fully formed α-dystroglycan due to the silencing of like-acetylglucosaminyltransferase (LARGE), in conjunction with altered integrin expression and regulated ECM degradation, have been reported to contribute to the increased metastatic potential of epithelial cells.^75^

Finally, S-linked GlcNAcylation is a recently discovered type of glycosylation, and has been reported on a limited number of proteins including Bassoon presynaptic cytomatrix protein. This protein was found in bacterial glycopeptides^76^ and mouse and rat synaptosome.^76^ Our proteomic data reveals the presence of unmodified Bassoon peptides in almost all samples from different regions, but our glycoproteomics analysis demonstrates that the S-GlcNAc glycopeptide was primarily detected in benign regions, and was rarely detected in tumor or necrotic regions. The canine peptide sequence is conserved across mammalia and shares 94% similarity with the human sequence. Bassoon is present in the ribbon synapse and acts as a scaffold that anchors the synaptic ribbon and vesicles to the presynaptic cytomatrix. Recent reports suggest that Bassoon has other functions in the cytomatrix and may contribute to synaptic plasticity. The third coiled coil (CC3) of bassoon binds to CtBP1, a transcriptional co-repressor that can be shuttled from the presynaptic compartment to the nucleus during increased neuronal activity. The S-linked glycopeptide LDFGQGAGSPVcLAQVK is located in CC3. Mediation of transcription by glycosylation is a well-known form of epigenetic regulation; thus, it would be interesting to investigate whether S-linked glycans are utilized for the same function in cancer. If so, they may serve as more selective targets for treatment due to the rarity of S-GlcNAc expression.

Here we demonstrated the utility of spatially-resolved glycoproteomics in complementing MALDI-MSI N-glycan imaging. We first showed that PNGAseF released N-glycans have differential expression in benign versus cancerous regions using MALDI-MSI on canine glioma FFPE tissues. We then subjected the separate regions to microdigestion followed by LC-MS/MS and exhaustive data analysis to show that: (a) intact N-glycopeptides linked MALDI MSI experiments to the associated glycoprotein, (b) a wide range of glycopeptide modifications could be identified, and (c) we could distinguish statistically significant glycan changes in benign, tumor, and necrotic regions. We note that this proof-of-principle experiment can be applied to any other FFPE bio-banked samples to identify diagnostic or prognostic indicators of various diseases. We envision that our MALDI MSI N-glycan imaging and spatially-resolved glycoproteomics workflow will find use in identifying targets for therapeutic intervention.

## Supporting information

Supplemental Data 1_Clinical glioma data

Supplemental Data 2_Proteomic pathway analysis

Supplemental data 3_Biantennary sialylated glycans

Supplemental Data 4_MSn Assignments

Supplemental Data 5_Total glycoforms detected

Supplemental Data 6_Lectin Staining

Supplemental Data 7_Glyccopeptides by sample

Supplemental Data 8_Glycopeptides by glycan type

## Author contributions

S.A.M., J.Q., and M.S. conceived the project. S.A.M., J.Q., A.R.R, F.K., S.A., and S.T. performed experiments and analyzed data. C.R.B., I.F., M.S. advised all mentees. S.A.M. and J.Q. wrote the manuscript with input from all authors.

## Notes

### Competing Interest Statement

C.R.B. is a cofounder and Scientific Advisory Board member of Lycia Therapeutics, Palleon Pharmaceuticals, Enable Bioscience, Redwood Biosciences (a subsidiary of Catalent), and InterVenn Biosciences, and a member of the Board of Directors of Eli Lilly & Company.

## References Cited

1. Koehler, J. W. et al. A Revised Diagnostic Classification of Canine Glioma: Towards Validation of the Canine Glioma Patient as a Naturally Occurring Preclinical Model for Human Glioma. Journal of Neuropathology & Experimental Neurology 77, 1039–1054 (2018).

2. Bentley, R. T., Ahmed, A. U., Yanke, A. B., Cohen-Gadol, A. A. & Dey, M. Dogs are man’s best friend: in sickness and in health. Neuro Oncol now 109 (2016) doi:10.1093/neuonc/now109.

3. Hubbard, M. E. et al. Naturally Occurring Canine Glioma as a Model for Novel Therapeutics. Cancer Investigation 36, 415–423 (2018).

4. Mitchell, D. et al. Common Molecular Alterations in Canine Oligodendroglioma and Human Malignant Gliomas and Potential Novel Therapeutic Targets. Front. Oncol. 9, 780 (2019).

5. Dickinson, P. J. Advances in Diagnostic and Treatment Modalities for Intracranial Tumors. J Vet Intern Med 28, 1165–1185 (2014).

6. Moremen, K. W., Tiemeyer, M. & Nairn, A. V. Vertebrate protein glycosylation: diversity, synthesis and function. Nat Rev Mol Cell Biol 13, 448–462 (2012).

7. Rudd, P., Karlsson, N. G., Khoo, K.-H. & Packer, N. H. Glycomics and Glycoproteomics. in Essentials of Glycobiology (eds. Varki, A. et al.) (Cold Spring Harbor Laboratory Press, 2015).

8. Pinho, S. S. & Reis, C. A. Glycosylation in cancer: mechanisms and clinical implications. Nat Rev Cancer 15, 540–555 (2015).

9. Kufe, D. W. Mucins in cancer: function, prognosis and therapy. Nat Rev Cancer 9, 874–885 (2009).

10. Hart, G. W. & Akimoto, Y. The O-GlcNAc Modification. in Essentials of Glycobiology (eds. Varki, A. et al.) (Cold Spring Harbor Laboratory Press, 2009).

11. Khidekel, N., Ficarro, S. B., Peters, E. C. & Hsieh-Wilson, L. C. Exploring the O-GlcNAc proteome: Direct identification of O-GlcNAc-modified proteins from the brain. Proceedings of the National Academy of Sciences 101, 13132–13137 (2004).

12. Vester-Christensen, M. B. et al. Mining the O-mannose glycoproteome reveals cadherins as major O-mannosylated glycoproteins. Proceedings of the National Academy of Sciences 110, 21018–21023 (2013).

13. Sheikh, M. O., Halmo, S. M. & Wells, L. Recent advancements in understanding mammalian O-mannosylation. Glycobiology 27, 806–819 (2017).

14. Larsen, I. S. B. et al. Mammalian *O* - mannosylation of cadherins and plexins is independent of protein *O* - mannosyltransferases 1 and 2. J. Biol. Chem. 292, 11586–11598 (2017).

15. Larsen, I. S. B., Narimatsu, Y., Clausen, H., Joshi, H. J. & Halim, A. Multiple distinct O-Mannosylation pathways in eukaryotes. Current Opinion in Structural Biology 56, 171–178 (2019).

16. Dobson, C. M., Hempel, S. J., Stalnaker, S. H., Stuart, R. & Wells, L. O-Mannosylation and human disease. Cell. Mol. Life Sci. 70, 2849–2857 (2013).

17. Moloney, D. J. et al. Mammalian Notch1 Is Modified with Two Unusual Forms of *O* - Linked Glycosylation Found on Epidermal Growth Factor-like Modules. J. Biol. Chem. 275, 9604–9611 (2000).

18. Li, Z. et al. Structural basis of Notch O-glucosylation and O–xylosylation by mammalian protein–O-glucosyltransferase 1 (POGLUT1). Nat Commun 8, 185 (2017).

19. Darula, Z. & Medzihradszky, K. F. Analysis of Mammalian O-Glycopeptides—We Have Made a Good Start, but There is a Long Way to Go. Mol Cell Proteomics 17, 2–17 (2018).

20. Leney, A. C., El Atmioui, D., Wu, W., Ovaa, H. & Heck, A. J. R. Elucidating crosstalk mechanisms between phosphorylation and O-GlcNAcylation. Proc Natl Acad Sci USA 114, E7255–E7261 (2017).

21. Hart, G. W., Slawson, C., Ramirez-Correa, G. & Lagerlof, O. Cross Talk Between O-GlcNAcylation and Phosphorylation: Roles in Signaling, Transcription, and Chronic Disease. Annu. Rev. Biochem. 80, 825–858 (2011).

22. Maynard, J. C., Burlingame, A. L. & Medzihradszky, K. F. Cysteine S-linked *N* - acetylglucosamine (S-GlcNAcylation), A New Post-translational Modification in Mammals. Mol Cell Proteomics 15, 3405–3411 (2016).

23. Veillon, L., Fakih, C., Abou-El-Hassan, H., Kobeissy, F. & Mechref, Y. Glycosylation Changes in Brain Cancer. ACS Chem. Neurosci. 9, 51–72 (2018).

24. Bull, C., Stoel, M. A., den Brok, M. H. & Adema, G. J. Sialic Acids Sweeten a Tumor’s Life. Cancer Research 74, 3199–3204 (2014).

25. Pearce, O. M. T. & Läubli, H. Sialic acids in cancer biology and immunity. Glycobiology 26, 111–128 (2016).

26. Varki, A., Kannagi, R., Toole, B. & Stanley, P. Glycosylation Changes in Cancer. in Essentials of Glycobiology (eds. Varki, A. et al.) (Cold Spring Harbor Laboratory Press, 2015).

27. Tachibana, H. et al. Changes of monosaccharide availability of human hybridoma lead to alteration of biological properties of human monoclonal antibody. Cytotechnology 16, 151–157 (1994).

28. Hanes, M. S., Moremen, K. W. & Cummings, R. D. Biochemical characterization of functional domains of the chaperone Cosmc. PLoS ONE 12, e0180242 (2017).

29. Dennis, J., Laferte, S., Waghorne, C., Breitman, M. & Kerbel, R. Beta 1-6 branching of Asn-linked oligosaccharides is directly associated with metastasis. Science 236, 582–585 (1987).

30. Noda, K. et al. Gene expression of?1-6 fucosyltransferase in human hepatoma tissues: A possible implication for increased fucosylation of ?-fetoprotein. Hepatology 28, 944–952 (1998).

31. Büll, C. et al. Sialic acid blockade suppresses tumor growth by enhancing T cell-mediated tumor immunity. Cancer Res canres. 3376. 2017 (2018) doi:10.1158/0008-5472.CAN-17-3376.

32. Thaysen-Andersen, M., Packer, N. H. & Schulz, B. L. Maturing Glycoproteomics Technologies Provide Unique Structural Insights into the *N* - glycoproteome and Its Regulation in Health and Disease. Mol Cell Proteomics 15, 1773–1790 (2016).

33. Furukawa, J. et al. Comprehensive Glycomics of a Multistep Human Brain Tumor Model Reveals Specific Glycosylation Patterns Related to Malignancy. PLoS ONE 10, e0128300 (2015).

34. Kudelka, M. R., Stowell, S. R., Cummings, R. D. & Neish, A. S. Intestinal epithelial glycosylation in homeostasis and gut microbiota interactions in IBD. Nat Rev Gastroenterol Hepatol (2020) doi:10.1038/s41575-020-0331-7.

35. Varki, A. & Gagneux, P. Multifarious roles of sialic acids in immunity: Roles of sialic acids in immunity. Annals of the New York Academy of Sciences 1253, 16–36 (2012).

36. Blaum, B. S. et al. Structural basis for sialic acid–mediated self-recognition by complement factor H. Nat Chem Biol 11, 77–82 (2015).

37. Hudak, J. E., Canham, S. M. & Bertozzi, C. R. Glycocalyx engineering reveals a Siglec-based mechanism for NK cell immunoevasion. Nat Chem Biol 10, 69–75 (2014).

38. Macauley, M. S., Crocker, P. R. & Paulson, J. C. Siglec-mediated regulation of immune cell function in disease. Nat Rev Immunol 14, 653–666 (2014).

39. Varki, A. Sialic acids in human health and disease. Trends in Molecular Medicine 14, 351–360 (2008).

40. Gray, M. A. et al. Targeted glycan degradation potentiates the anticancer immune response in vivo. Nat Chem Biol (2020) doi:10.1038/s41589-020-0622-x.

41. Xiao, H., Woods, E. C., Vukojicic, P. & Bertozzi, C. R. Precision glycocalyx editing as a strategy for cancer immunotherapy. Proc Natl Acad Sci USA 113, 10304–10309 (2016).

42. Posey, A. D. et al. Engineered CAR T Cells Targeting the Cancer-Associated Tn-Glycoform of the Membrane Mucin MUC1 Control Adenocarcinoma. Immunity 44, 1444–1454 (2016).

43. Olsen, J. V. et al. Higher-energy C-trap dissociation for peptide modification analysis. Nat Methods 4, 709–712 (2007).

44. Syka, J. E. P., Coon, J. J., Schroeder, M. J., Shabanowitz, J. & Hunt, D. F. Peptide and protein sequence analysis by electron transfer dissociation mass spectrometry. Proceedings of the National Academy of Sciences 101, 9528–9533 (2004).

45. Reiding, K. R., Bondt, A., Franc, V. & Heck, A. J. R. The benefits of hybrid fragmentation methods for glycoproteomics. TrAC Trends in Analytical Chemistry 108, 260–268 (2018).

46. Riley, N. M., Westphall, M. S. & Coon, J. J. Activated Ion-Electron Transfer Dissociation Enables Comprehensive Top-Down Protein Fragmentation. J. Proteome Res. 16, 2653–2659 (2017).

47. Shajahan, A., Supekar, N. T., Gleinich, A. S. & Azadi, P. Deducing the N-and O-glycosylation profile of the spike protein of novel coronavirus SARS-CoV-2. Glycobiology cwaa042 (2020) doi:10.1093/glycob/cwaa042.

48. Riley, N. M., Malaker, S. A., Driessen, M. & Bertozzi, C. R. Optimal Dissociation Methods Differ for N-and O-glycopeptides. J. Proteome Res. acs.jproteome.0c00218 (2020) doi:10.1021/acs.jproteome.0c00218.

49. Drake, R. R. et al. MALDI Mass Spectrometry Imaging of N-Linked Glycans in Cancer Tissues. in Advances in Cancer Research vol. 134 85–116 (Elsevier, 2017).

50. Briggs, M. T. et al. MALDI mass spectrometry imaging of *N*-glycans on tibial cartilage and subchondral bone proteins in knee osteoarthritis. Proteomics 16, 1736–1741 (2016).

51. Everest-Dass, A. V. et al. N-glycan MALDI Imaging Mass Spectrometry on Formalin-Fixed Paraffin-Embedded Tissue Enables the Delineation of Ovarian Cancer Tissues. Mol Cell Proteomics 15, 3003–3016 (2016).

52. Powers, T. W. et al. MALDI Imaging Mass Spectrometry Profiling of N-Glycans in Formalin-Fixed Paraffin Embedded Clinical Tissue Blocks and Tissue Microarrays. PLoS ONE 9, e106255 (2014).

53. Powers, T., Holst, S., Wuhrer, M., Mehta, A. & Drake, R. Two-Dimensional N-Glycan Distribution Mapping of Hepatocellular Carcinoma Tissues by MALDI-Imaging Mass Spectrometry. Biomolecules 5, 2554–2572 (2015).

54. Holst, S. et al. Linkage-Specific *in Situ* Sialic Acid Derivatization for N-Glycan Mass Spectrometry Imaging of Formalin-Fixed Paraffin-Embedded Tissues. Anal. Chem. 88, 5904–5913 (2016).

55. Heijs, B. et al. Multimodal Mass Spectrometry Imaging of *N* - Glycans and Proteins from the Same Tissue Section. Anal. Chem. 88, 7745–7753 (2016).

56. Angel, P. M., Mehta, A., Norris-Caneda, K. & Drake, R. R. MALDI Imaging Mass Spectrometry of N-glycans and Tryptic Peptides from the Same Formalin-Fixed, Paraffin-Embedded Tissue Section. in Tissue Proteomics (eds. Sarwal, M. M. & Sigdel, T. K.) vol. 1788 225–241 (Springer New York, 2017).

57. Angel, P. M. et al. Mapping Extracellular Matrix Proteins in Formalin-Fixed, Paraffin-Embedded Tissues by MALDI Imaging Mass Spectrometry. J. Proteome Res. 17, 635–646 (2018).

58. Quanico, J. et al. Development of liquid microjunction extraction strategy for improving protein identification from tissue sections. Journal of Proteomics 79, 200–218 (2013).

59. Wisztorski, M. et al. Microproteomics by liquid extraction surface analysis: Application to FFPE tissue to study the fimbria region of tubo-ovarian cancer. Prot. Clin. Appl. 7, 234–240 (2013).

60. Quanico, J. et al. NanoLC-MS coupling of liquid microjunction microextraction for on-tissue proteomic analysis. Biochimica et Biophysica Acta (BBA) - Proteins and Proteomics 1865, 891–900 (2017).

61. Wisztorski, M. et al. Droplet-Based Liquid Extraction for Spatially-Resolved Microproteomics Analysis of Tissue Sections. in Imaging Mass Spectrometry (ed. Cole, L. M.) vol. 1618 49–63 (Springer New York, 2017).

62. Yuryev, A., Kotelnikova, E. & Daraselia, N. Ariadne’s ChemEffect and Pathway Studio knowledge base. Expert Opinion on Drug Discovery 4, 1307–1318 (2009).

63. Bonnet, A. et al. Pathway results from the chicken data set using GOTM, Pathway Studio and Ingenuity softwares. BMC Proc 3, S11 (2009).

64. Kobeissy, F. H. et al. Neuroproteomics and Systems Biology Approach to Identify Temporal Biomarker Changes Post Experimental Traumatic Brain Injury in Rats. Front. Neurol. 7, (2016).

65. Heberle, H., Meirelles, G. V., da Silva, F. R., Telles, G. P. & Minghim, R. InteractiVenn: a web-based tool for the analysis of sets through Venn diagrams. BMC Bioinformatics 16, 169 (2015).

66. Dwyer, C. A., Baker, E., Hu, H. & Matthews, R. T. RPTPζ/phosphacan is abnormally glycosylated in a model of muscle–eye–brain disease lacking functional POMGnT1. Neuroscience 220, 47–61 (2012).

67. Rutherford, M. A. & Pangršič, T. Molecular anatomy and physiology of exocytosis in sensory hair cells. Cell Calcium 52, 327–337 (2012).

68. Zhang, S., Shang, S., Li, W., Qin, X. & Liu, Y. Insights on N-glycosylation of human haptoglobin and its association with cancers. Glycobiology 26, 684–692 (2016).

69. Fujimura, T. et al. Glycosylation status of haptoglobin in sera of patients with prostate cancervs. benign prostate disease or normal subjects. Int. J. Cancer 122, 39–49 (2008).

70. Lu, R. et al. The role of brevican in glioma: promoting tumor cell motility in vitro and in vivo. BMC Cancer 12, 607 (2012).

71. Hu, B., Kong, L. L., Matthews, R. T. & Viapiano, M. S. The Proteoglycan Brevican Binds to Fibronectin after Proteolytic Cleavage and Promotes Glioma Cell Motility. J. Biol. Chem. 283, 24848–24859 (2008).

72. Viapiano, M. S., Bi, W. L., Piepmeier, J., Hockfield, S. & Matthews, R. T. Novel Tumor-Specific Isoforms of BEHAB/Brevican Identified in Human Malignant Gliomas. Cancer Res 65, 6726–6733 (2005).

73. Sim, H., Hu, B. & Viapiano, M. S. Reduced Expression of the Hyaluronan and Proteoglycan Link Proteins in Malignant Gliomas. J. Biol. Chem. 284, 26547–26556 (2009).

74. Steentoft, C. et al. Precision mapping of the human O-GalNAc glycoproteome through SimpleCell technology. EMBO J. 32, 1478–1488 (2013).

75. de Bernabé, D. B.-V. et al. Loss of α-Dystroglycan Laminin Binding in Epithelium-derived Cancers Is Caused by Silencing of *LARGE*. J. Biol. Chem. 284, 11279–11284 (2009).

76. Stepper, J. et al. Cysteine *S* - glycosylation, a new post-translational modification found in glycopeptide bacteriocins. FEBS Letters 585, 645–650 (2011).

