## Supplemental Data 1_Clinical glioma data for "On-tissue spatially-resolved glycoproteomics guided by N-glycan imaging reveal global dysregulation of canine glioma glycoproteomic landscape"

### Supplemental Data 1: Clinical data of Glioma Samples

| Code | Breed | Age (years) | Sex | N/US | Type of Cancer | Area of brain |
|---|---|---|---|---|---|---|
| VH 15 1139A<br>VH 15 1139D | Boxer | 9 | F | Neutered | Glioblastoma (Grade IV WHO) | Cortex left hemisphere |
| VH 16 0440C<br>VH 16 0440D | French bulldog | 7 | M | Unspayed | Anaplastic Oligodendroglioma (Grade III WHO) | Cortex left hemisphere |
| VH 16 0703A<br>VH 16 0703B | French bulldog | 8 | M | Unspayed | Anaplastic Oligodendroglioma (Grade III WHO) | Cortex Right hemisphere |
| VH 13 0935 | French bulldog | 8 | F | Neutered | Anaplastic Oligodendroglioma (Grade III WHO) | Left parietal lobe (cranial aspect) |
| VH 14 0622 | Border Collie | 13 | F | Unspayed | Glioblastoma (Grade IV WHO) | junction between parietal and frontal lobe left hemisphere |
| VH 15 3520A | English bulldog | 5 | F | Unspayed | Anaplastic Oligodendroglioma (Grade III WHO) | Left rhinencephalus |
