## Supplementary figures and images for "On-tissue spatially-resolved glycoproteomics guided by N-glycan imaging reveal global dysregulation of canine glioma glycoproteomic landscape"

### Supplemental data 3_Biantennary sialylated glycans

#### Supplementary Data 3. Ion traces of biantennary sialylated N-glycans

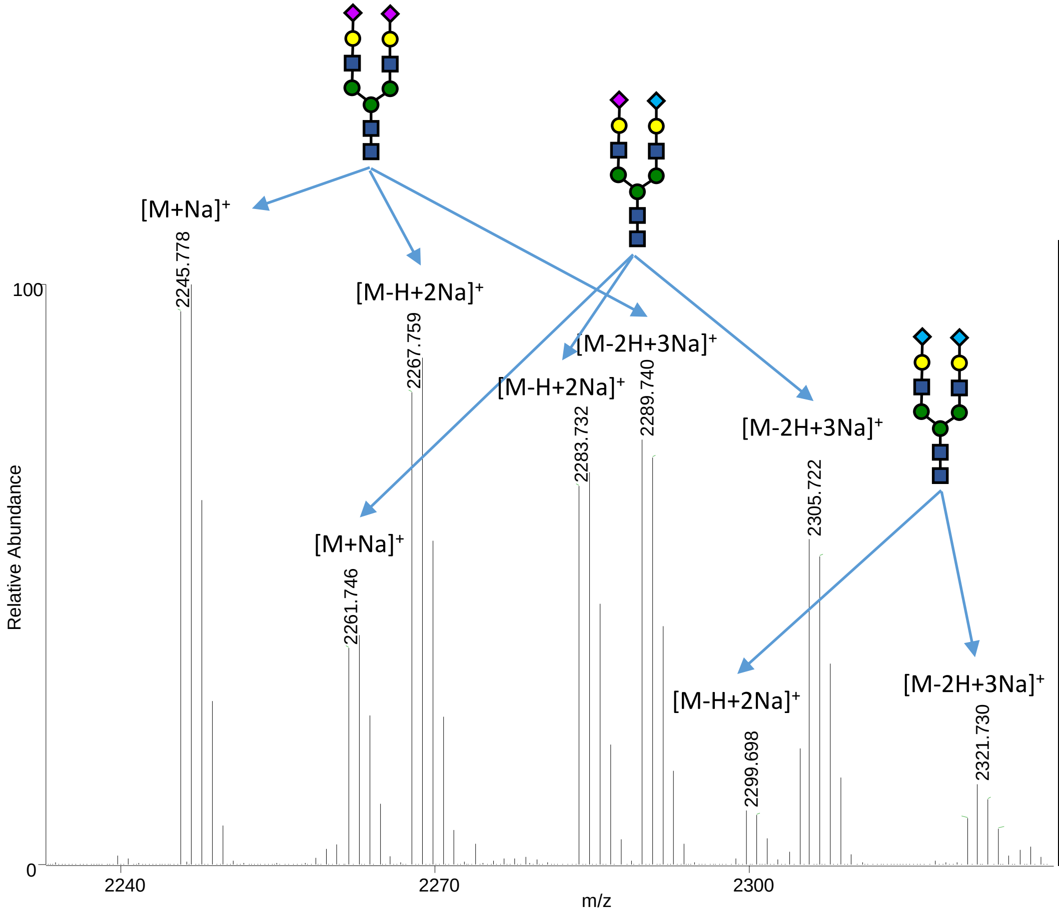


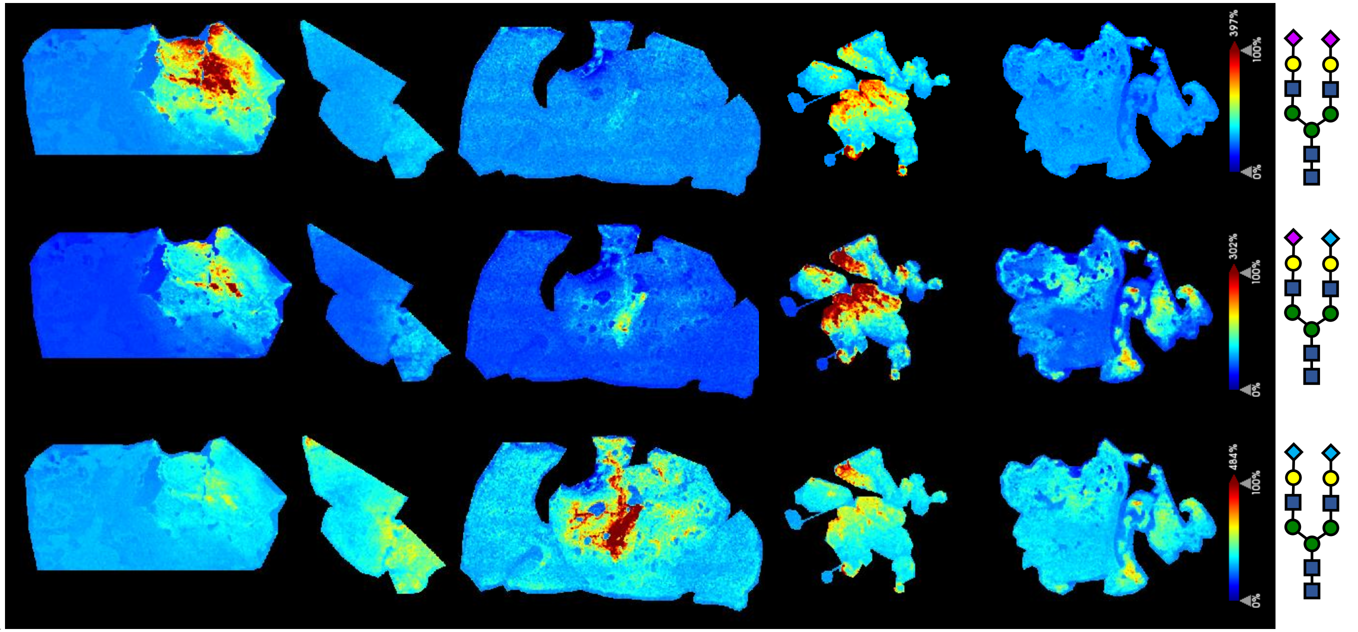
