## Supplemental Data 4_MSn Assignments for "On-tissue spatially-resolved glycoproteomics guided by N-glycan imaging reveal global dysregulation of canine glioma glycoproteomic landscape"

**Supplementary Data 4. MSn assignments.** MSn spectra were acquired directly on tissue using a MALDI LTQ Orbitrap instrument after images had been obtained. The MALDI LTQ Orbitrap XL is equipped with a commercial N2 laser (LTB Lasertechnik, Berlin, Germany) operating at $\lambda = 337$ nm with a maximum repetition rate of 60 Hz. The hybrid configuration replaces the heated capillary of the electrospray source with a q00 that sends packets of ions into a linear trap for collision-induced fragmentation (CID), with the fragment ions then being concentrated in a C-trap and transferred to the orbitrap for high-resolution mass analysis. The maximum energy per pulse was set to 12 µJ. Precursor ion isolation was performed using an isolation window between ±1 and ±3 Da and the fragments scanned with a maximum accumulation time of 120 ms. Succeeding MSn of the daughter ions were performed with a maximum accumulation time of 180 ms. External calibration was performed using the ProteoMass MALDI Calibration Kit (Sigma-Aldrich, St. Quentin-Fallavier, France).

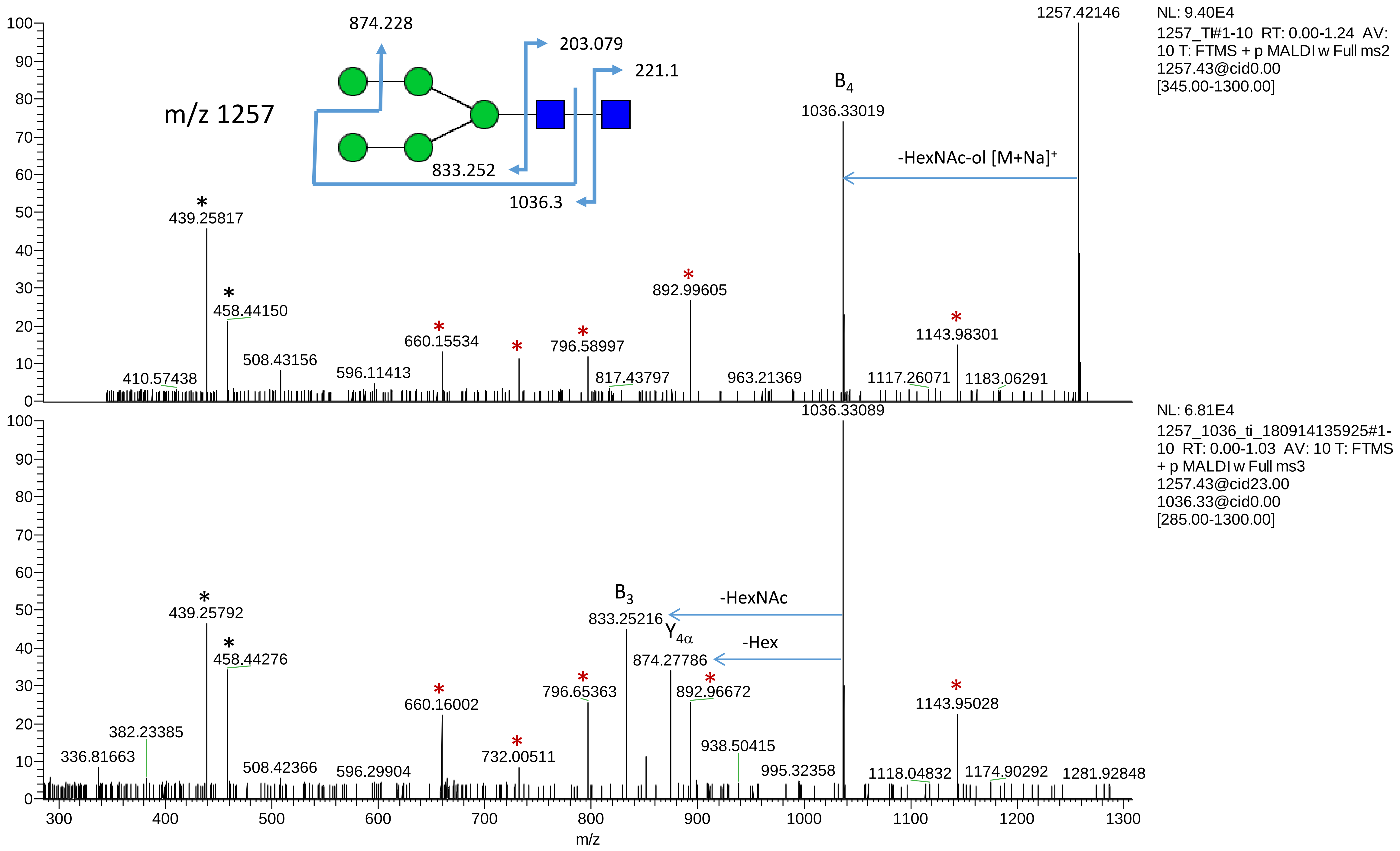
m/z 1257
874.228
203.079
221.1
833.252
1036.3
B4
1036.33019
-HexNAc-ol [M+Na]+
1257.42146
NL: 9.40E4
1257_TI#1-10 RT: 0.00-1.24 AV: 10 T: FTMS + p MALDI w Full ms2 1257.43@cid0.00 [345.00-1300.00]
439.25817
458.44150
508.43156
410.57438
596.11413
660.15534
796.58997
817.43797
892.99605
963.21369
1117.26071
1143.98301
1183.06291
1036.33089
NL: 6.81E4
1257_1036_ti_180914135925#1-10 RT: 0.00-1.03 AV: 10 T: FTMS + p MALDI w Full ms3 1257.43@cid23.00 1036.33@cid0.00 [285.00-1300.00]
B3
833.25216
-HexNAc
Y4α
874.27786
-Hex
439.25792
458.44276
382.23385
336.81663
508.42366
596.29904
660.16002
732.00511
796.65363
892.96672
938.50415
995.32358
1118.04832
1143.95028
1174.90292
1281.92848
m/z

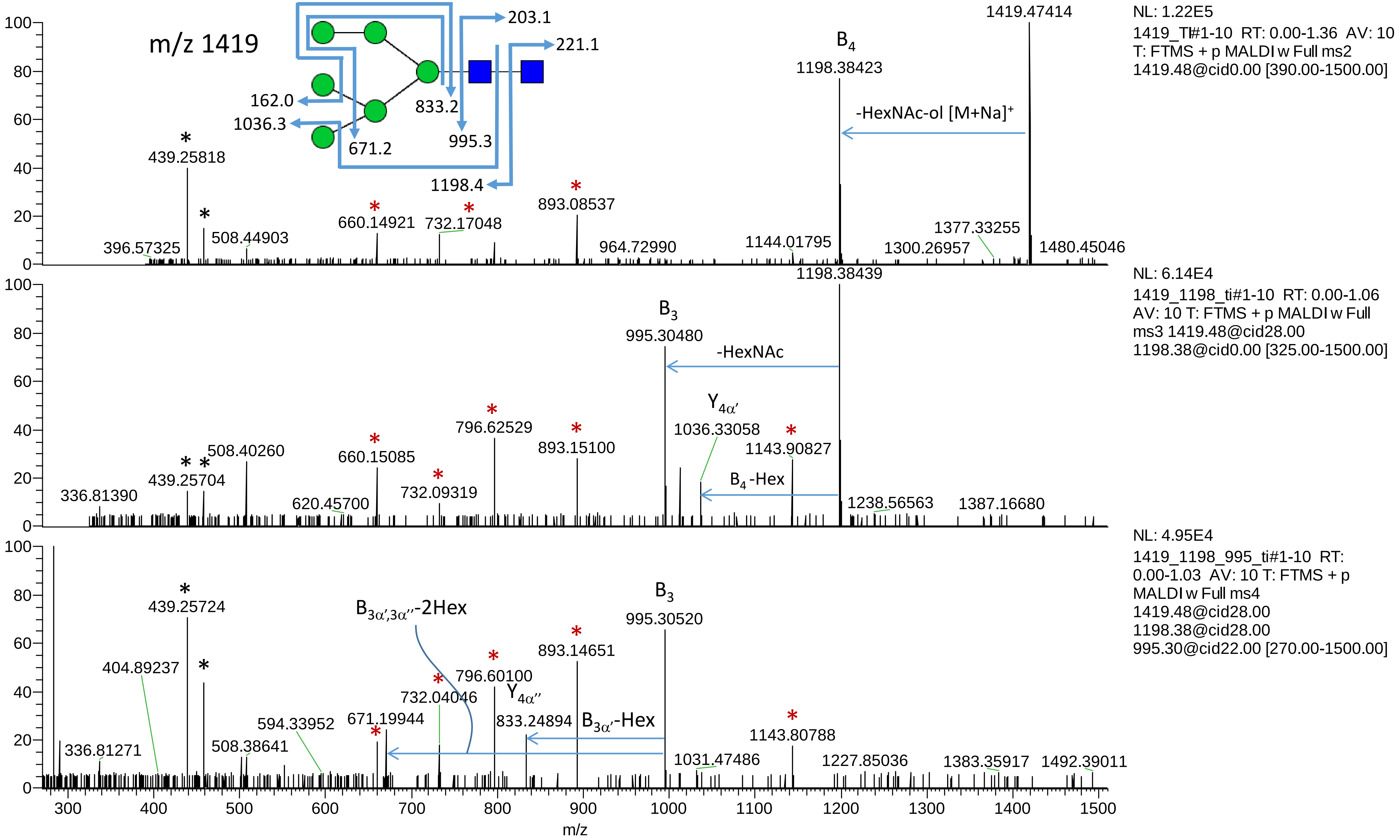
m/z 1419
203.1
221.1
162.0
1036.3
833.2
671.2
995.3
1198.4
1419.47414
B4
1198.38423
-HexNAc-ol [M+Na]+
439.25818
396.57325
508.44903
660.14921
732.17048
893.08537
964.72990
1144.01795
1300.26957
1377.33255
1480.45046
NL: 1.22E5
1419_TI#1-10 RT: 0.00-1.36 AV: 10
T: FTMS + p MALDI w Full ms2
1419.48@cid0.00 [390.00-1500.00]
1198.38439
B3
995.30480
-HexNAc
Y4α′
1036.33058
1143.90827
B4-Hex
336.81390
439.25704
508.40260
620.45700
660.15085
732.09319
796.62529
893.15100
1238.56563
1387.16680
NL: 6.14E4
1419_1198_ti#1-10 RT: 0.00-1.06
AV: 10 T: FTMS + p MALDI w Full
ms3 1419.48@cid28.00
1198.38@cid0.00 [325.00-1500.00]
439.25724
404.89237
336.81271
508.38641
594.33952
671.19944
B3α′,3α″-2Hex
732.04046
796.60100
Y4α″
833.24894
B3α′-Hex
893.14651
B3
995.30520
1031.47486
1143.80788
1227.85036
1383.35917
1492.39011
NL: 4.95E4
1419_1198_995_ti#1-10 RT:
0.00-1.03 AV: 10 T: FTMS + p
MALDI w Full ms4
1419.48@cid28.00
1198.38@cid28.00
995.30@cid22.00 [270.00-1500.00]
m/z

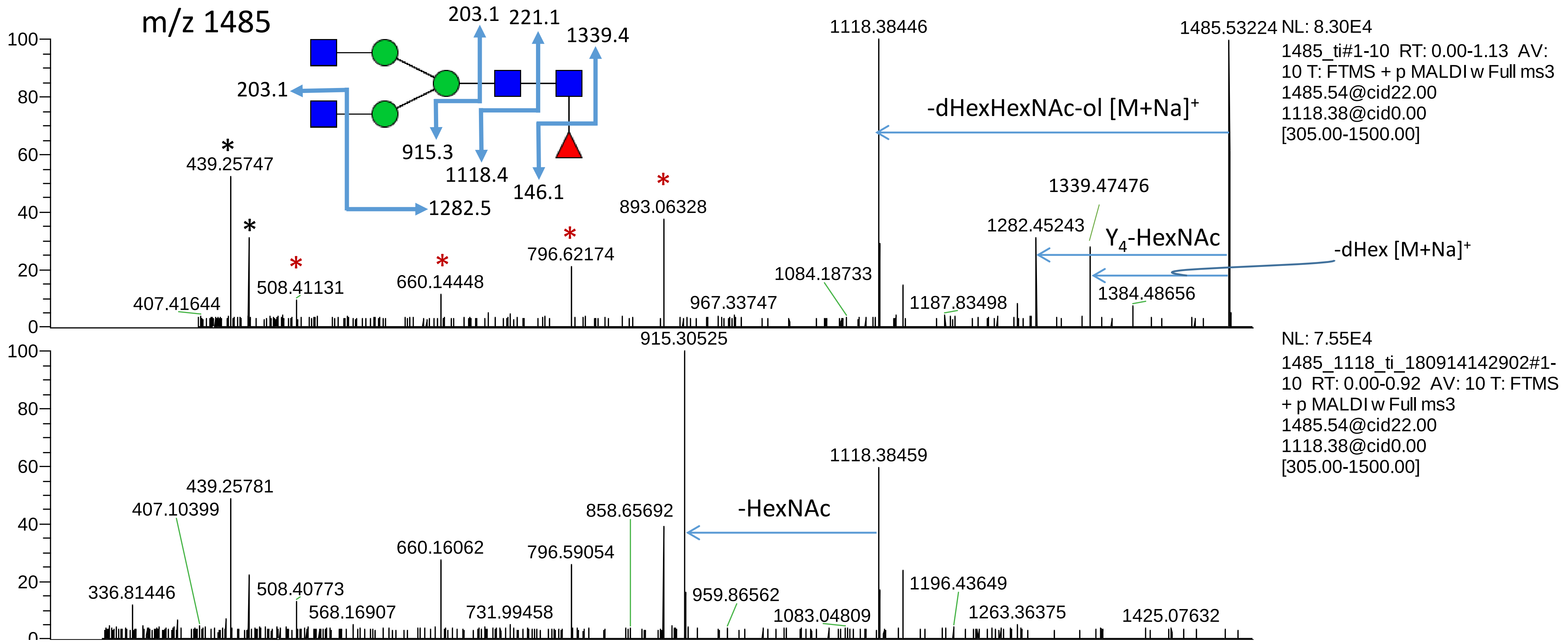
m/z 1485
203.1
221.1
1339.4
203.1
915.3
1118.4
146.1
1282.5
1118.38446
1485.53224 NL: 8.30E4
1485_ti#1-10 RT: 0.00-1.13 AV: 10 T: FTMS + p MALDI w Full ms3 1485.54@cid22.00 1118.38@cid0.00 [305.00-1500.00]
-dHexHexNAc-ol [M+Na]+
439.25747
893.06328
1339.47476
1282.45243
$Y_4$-HexNAc
-dHex [M+Na]+
508.41131
660.14448
796.62174
1084.18733
407.41644
967.33747
1187.83498
1384.48656
915.30525
NL: 7.55E4
1485_1118_ti_180914142902#1-10 RT: 0.00-0.92 AV: 10 T: FTMS + p MALDI w Full ms3 1485.54@cid22.00 1118.38@cid0.00 [305.00-1500.00]
1118.38459
439.25781
407.10399
858.65692
-HexNAc
660.16062
796.59054
336.81446
508.40773
568.16907
731.99458
959.86562
1083.04809
1196.43649
1263.36375
1425.07632

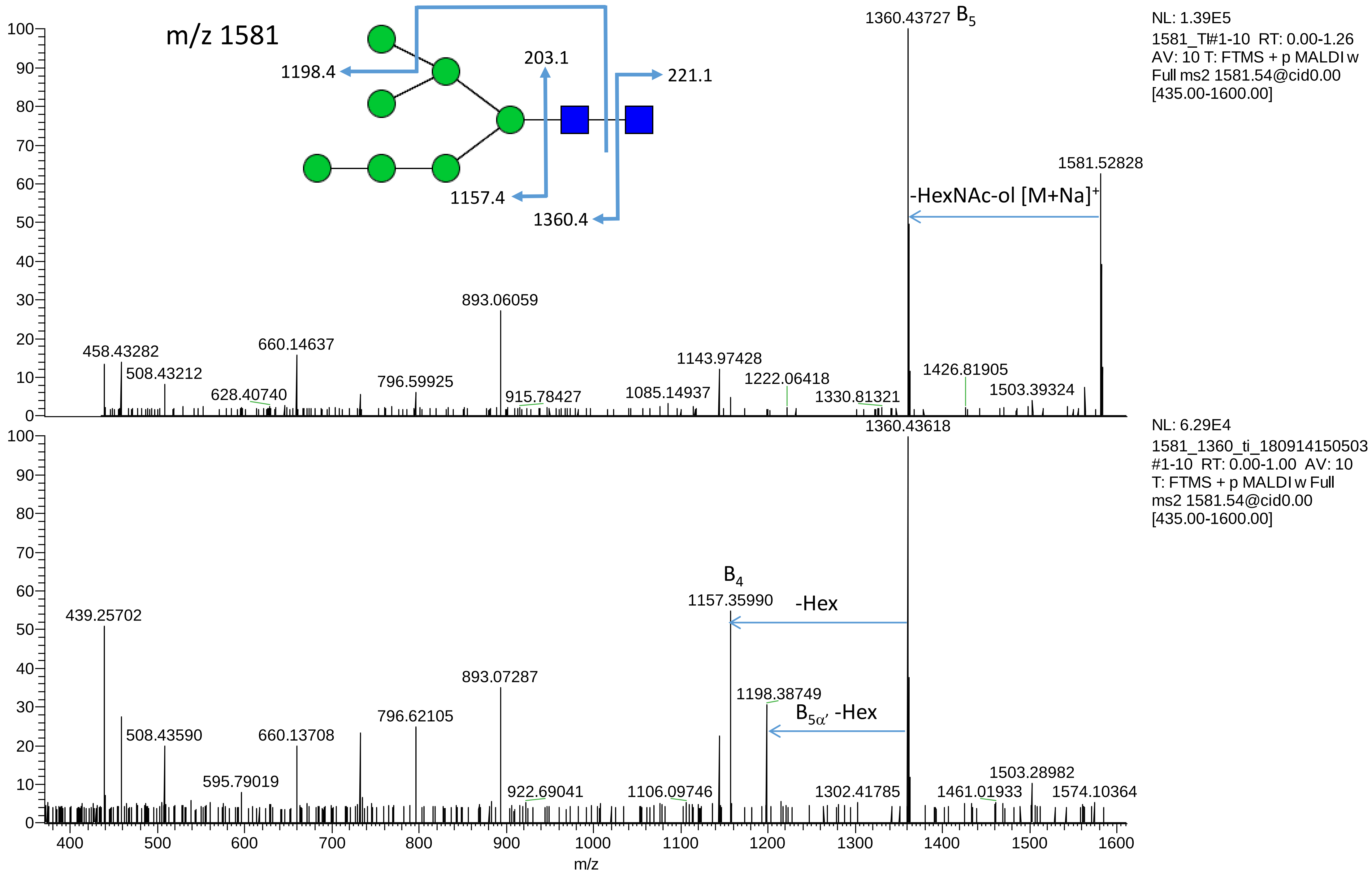

m/z 1581
1198.4
203.1
221.1
1157.4
1360.4
1360.43727 B5
NL: 1.39E5
1581_TI#1-10 RT: 0.00-1.26
AV: 10 T: FTMS + p MALDI w
Full ms2 1581.54@cid0.00
[435.00-1600.00]
1581.52828
-HexNAc-ol [M+Na]+
893.06059
458.43282
508.43212
660.14637
628.40740
796.59925
915.78427
1143.97428
1222.06418
1085.14937
1330.81321
1426.81905
1503.39324
1360.43618
NL: 6.29E4
1581_1360_ti_180914150503
#1-10 RT: 0.00-1.00 AV: 10
T: FTMS + p MALDI w Full
ms2 1581.54@cid0.00
[435.00-1600.00]
B4
1157.35990
-Hex
439.25702
893.07287
1198.38749
B5α′ -Hex
796.62105
508.43590
660.13708
595.79019
922.69041
1106.09746
1302.41785
1461.01933
1503.28982
1574.10364
m/z

m/z 1647

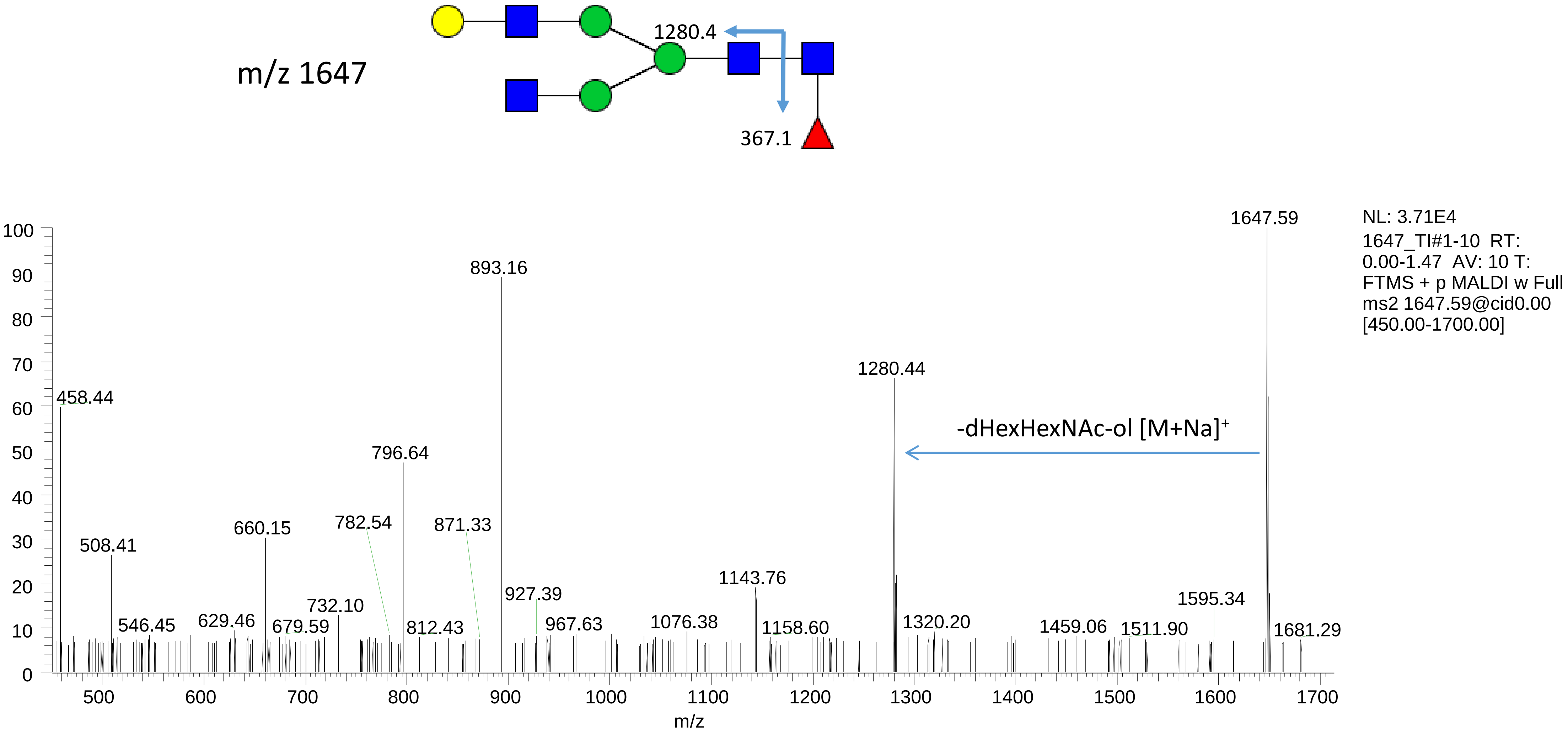
1280.4
367.1
NL: 3.71E4
1647_TI#1-10 RT: 0.00-1.47 AV: 10 T: FTMS + p MALDI w Full ms2 1647.59@cid0.00 [450.00-1700.00]
1647.59
893.16
1280.44
-dHexHexNAc-ol [M+Na]+
458.44
796.64
660.15
782.54
871.33
508.41
1143.76
927.39
1595.34
732.10
546.45
629.46
679.59
812.43
967.63
1076.38
1158.60
1320.20
1459.06
1511.90
1681.29
m/z

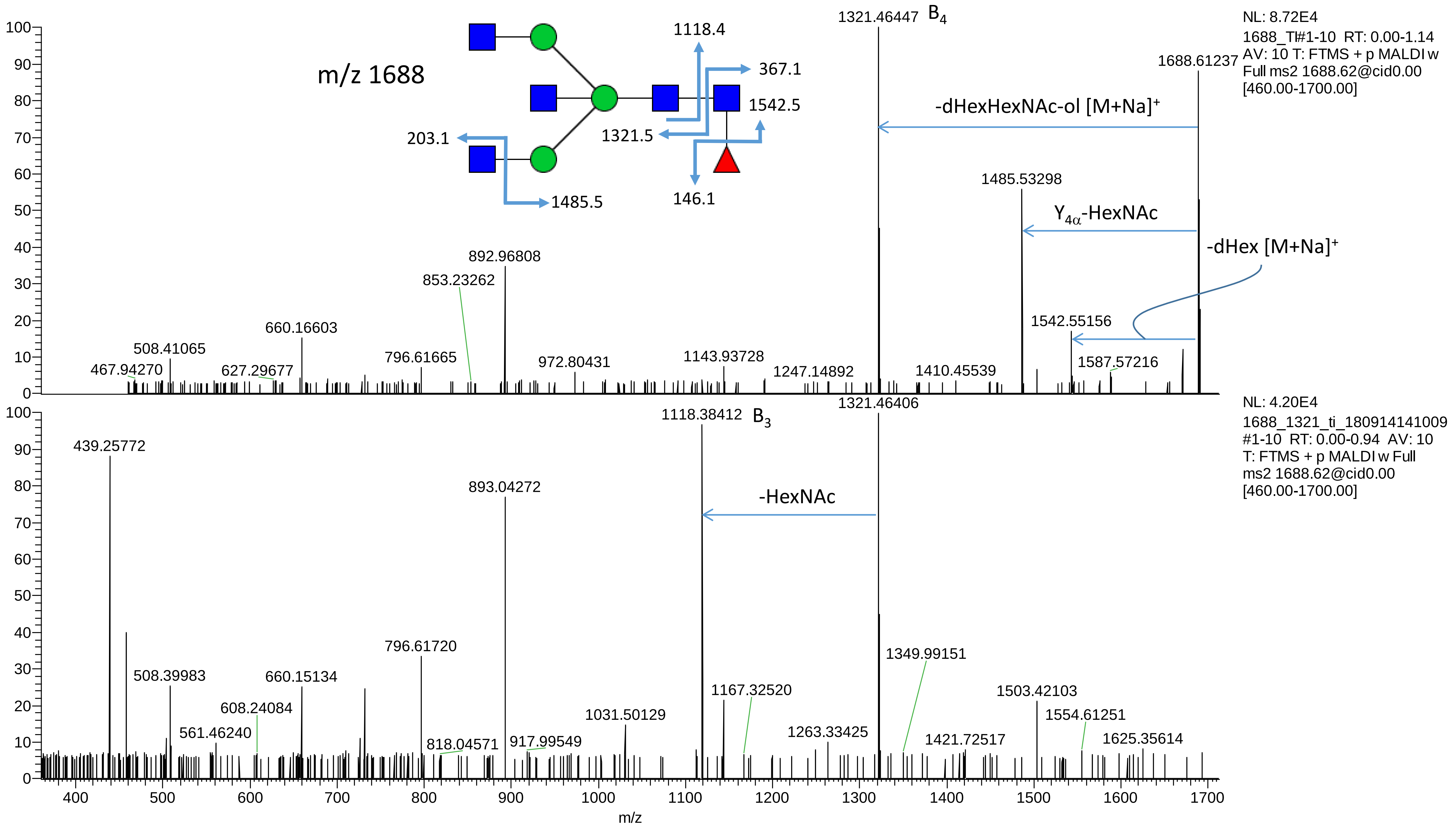

m/z 1688
1118.4
367.1
1542.5
1321.5
203.1
146.1
1485.5
1321.46447 B4
NL: 8.72E4
1688_TI#1-10 RT: 0.00-1.14
AV: 10 T: FTMS + p MALDI w
Full ms2 1688.62@cid0.00
[460.00-1700.00]
1688.61237
-dHexHexNAc-ol [M+Na]+
1485.53298
Y4α-HexNAc
-dHex [M+Na]+
892.96808
853.23262
660.16603
508.41065
467.94270
627.29677
796.61665
972.80431
1143.93728
1247.14892
1410.45539
1542.55156
1587.57216
NL: 4.20E4
1688_1321_ti_180914141009
#1-10 RT: 0.00-0.94 AV: 10
T: FTMS + p MALDI w Full
ms2 1688.62@cid0.00
[460.00-1700.00]
1118.38412 B3
1321.46406
-HexNAc
439.25772
893.04272
796.61720
508.39983
660.15134
608.24084
561.46240
818.04571
917.99549
1031.50129
1167.32520
1263.33425
1349.99151
1421.72517
1503.42103
1554.61251
1625.35614
m/z

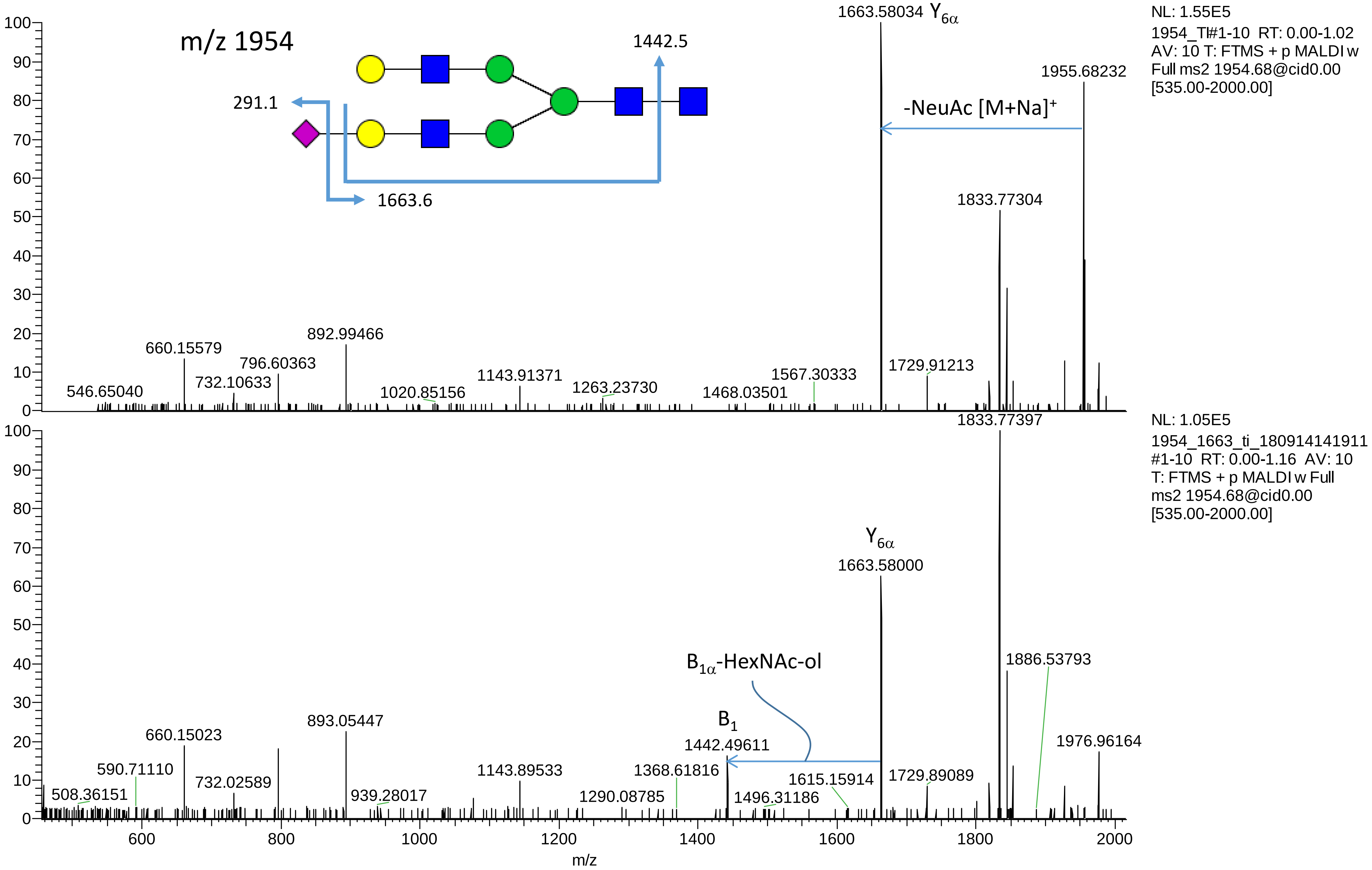

m/z 1954
291.1
1663.6
1442.5
1663.58034 Y6α
1955.68232
-NeuAc [M+Na]+
1833.77304
NL: 1.55E5
1954_TI#1-10 RT: 0.00-1.02
AV: 10 T: FTMS + p MALDI w
Full ms2 1954.68@cid0.00
[535.00-2000.00]
546.65040
660.15579
732.10633
796.60363
892.99466
1020.85156
1143.91371
1263.23730
1468.03501
1567.30333
1729.91213
1833.77397
NL: 1.05E5
1954_1663_ti_180914141911
#1-10 RT: 0.00-1.16 AV: 10
T: FTMS + p MALDI w Full
ms2 1954.68@cid0.00
[535.00-2000.00]
Y6α
1663.58000
B1α-HexNAc-ol
B1
1442.49611
1886.53793
1976.96164
508.36151
590.71110
660.15023
732.02589
893.05447
939.28017
1143.89533
1290.08785
1368.61816
1496.31186
1615.15914
1729.89089
m/z

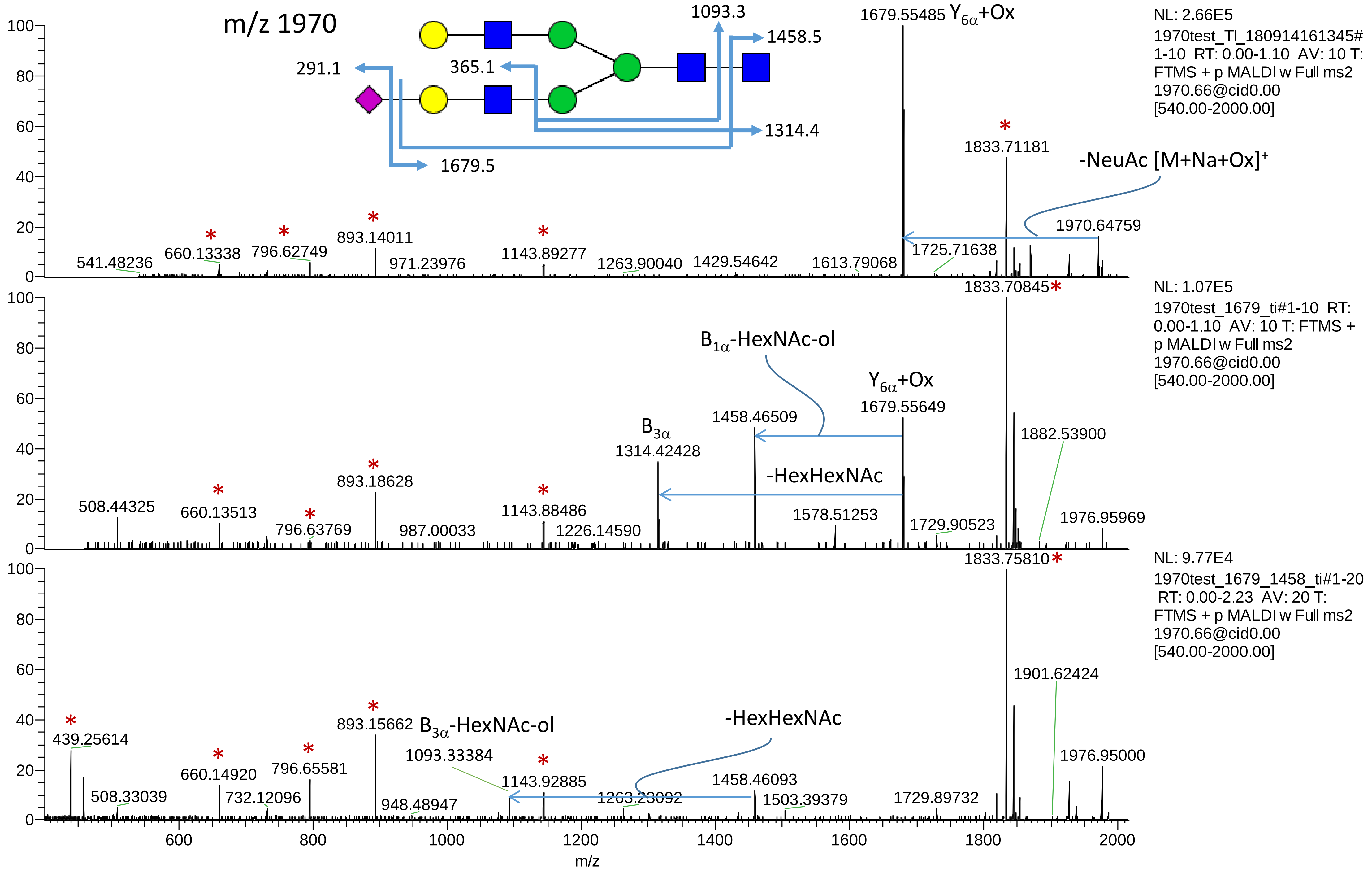
m/z 1970
1093.3
1458.5
291.1
365.1
1314.4
1679.5
1679.55485 Y6α+Ox
1833.71181
-NeuAc [M+Na+Ox]+
1970.64759
541.48236
660.13338
796.62749
893.14011
971.23976
1143.89277
1263.90040
1429.54642
1613.79068
1725.71638
NL: 2.66E5
1970test_TI_180914161345#
1-10 RT: 0.00-1.10 AV: 10 T:
FTMS + p MALDI w Full ms2
1970.66@cid0.00
[540.00-2000.00]
1833.70845
B1α-HexNAc-ol
Y6α+Ox
1679.55649
1458.46509
B3α
1314.42428
1882.53900
-HexHexNAc
508.44325
660.13513
796.63769
893.18628
987.00033
1143.88486
1226.14590
1578.51253
1729.90523
1976.95969
NL: 1.07E5
1970test_1679_ti#1-10 RT:
0.00-1.10 AV: 10 T: FTMS +
p MALDI w Full ms2
1970.66@cid0.00
[540.00-2000.00]
1833.75810
NL: 9.77E4
1970test_1679_1458_ti#1-20
RT: 0.00-2.23 AV: 20 T:
FTMS + p MALDI w Full ms2
1970.66@cid0.00
[540.00-2000.00]
1901.62424
439.25614
893.15662 B3α-HexNAc-ol
-HexHexNAc
1093.33384
1143.92885
660.14920
796.65581
508.33039
732.12096
948.48947
1263.23092
1458.46093
1503.39379
1729.89732
1976.95000
m/z

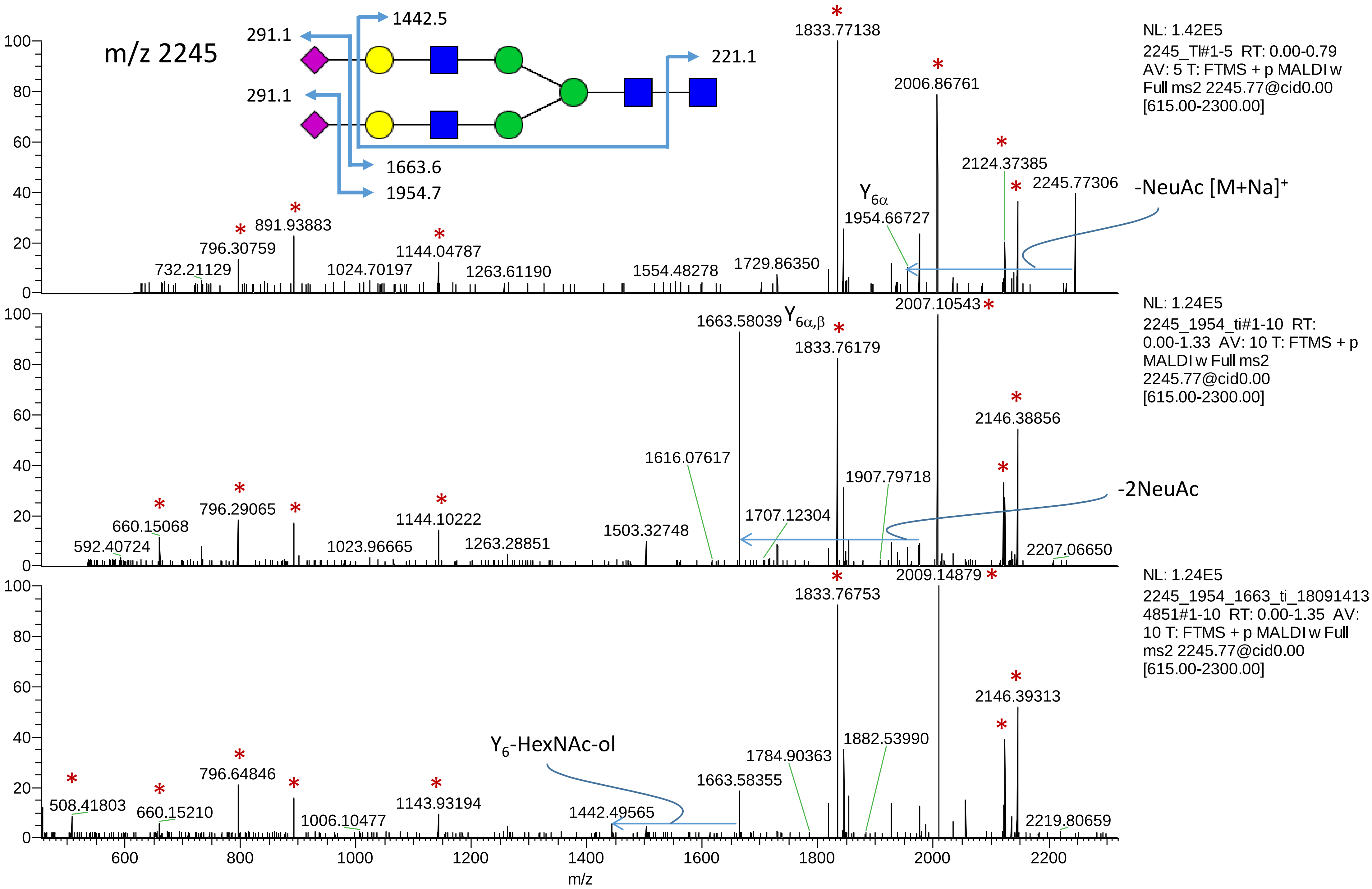
m/z 2245
291.1
291.1
1442.5
221.1
1663.6
1954.7
NL: 1.42E5
2245_TI#1-5 RT: 0.00-0.79
AV: 5 T: FTMS + p MALDI w
Full ms2 2245.77@cid0.00
[615.00-2300.00]
1833.77138
2006.86761
2124.37385
2245.77306
-NeuAc [M+Na]+
Y6α
1954.66727
891.93883
796.30759
732.21129
1024.70197
1144.04787
1263.61190
1554.48278
1729.86350
NL: 1.24E5
2245_1954_ti#1-10 RT:
0.00-1.33 AV: 10 T: FTMS + p
MALDI w Full ms2
2245.77@cid0.00
[615.00-2300.00]
2007.10543
1663.58039 Y6α,β
1833.76179
2146.38856
1616.07617
1907.79718
-2NeuAc
1707.12304
1503.32748
796.29065
660.15068
592.40724
1144.10222
1023.96665
1263.28851
2207.06650
NL: 1.24E5
2245_1954_1663_ti_18091413
4851#1-10 RT: 0.00-1.35 AV:
10 T: FTMS + p MALDI w Full
ms2 2245.77@cid0.00
[615.00-2300.00]
1833.76753
2009.14879
2146.39313
Y6-HexNAc-ol
1882.53990
1784.90363
1663.58355
796.64846
508.41803
660.15210
1143.93194
1006.10477
1442.49565
2219.80659
m/z

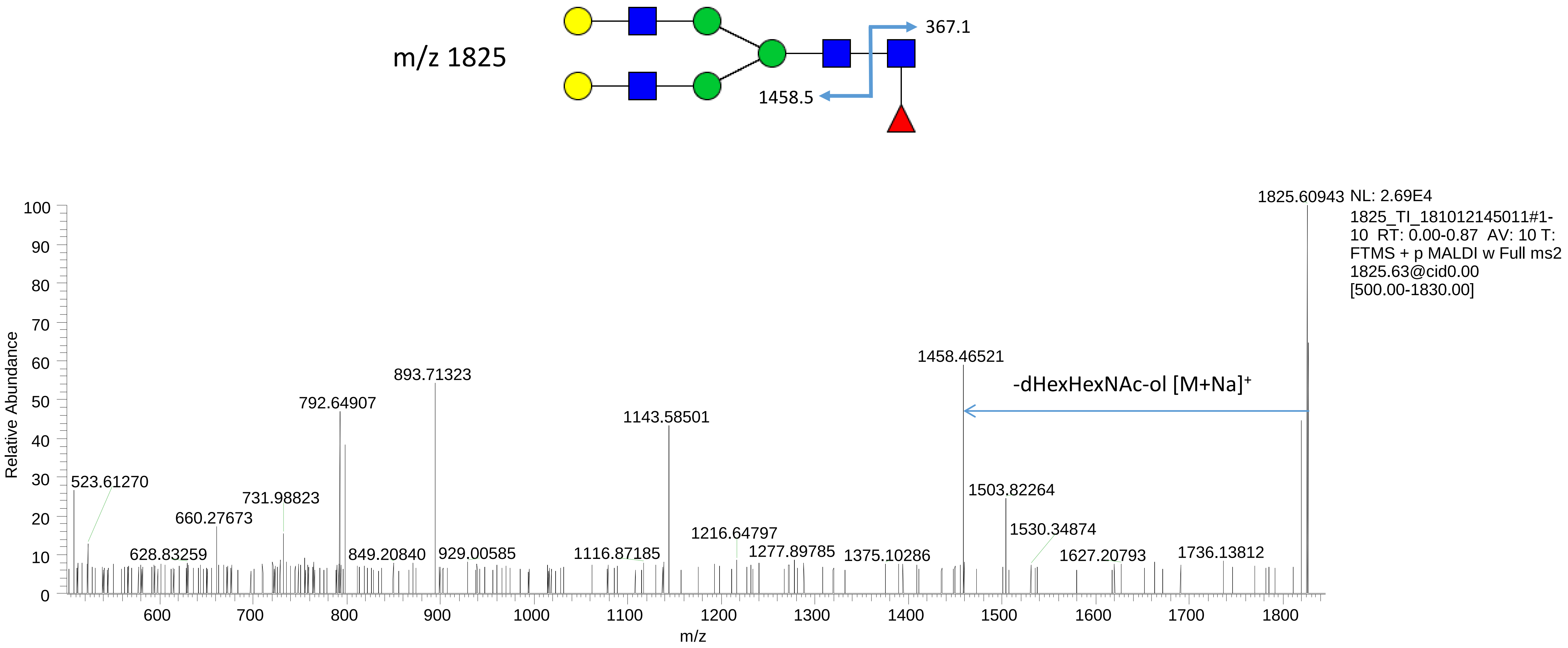
m/z 1825
367.1
1458.5
1825.60943
NL: 2.69E4
1825_TI_181012145011#1-10 RT: 0.00-0.87 AV: 10 T: FTMS + p MALDI w Full ms2 1825.63@cid0.00 [500.00-1830.00]
1458.46521
-dHexHexNAc-ol [M+Na]+
893.71323
792.64907
1143.58501
523.61270
731.98823
660.27673
1503.82264
1216.64797
1530.34874
628.83259
849.20840
929.00585
1116.87185
1277.89785
1375.10286
1627.20793
1736.13812
Relative Abundance
m/z

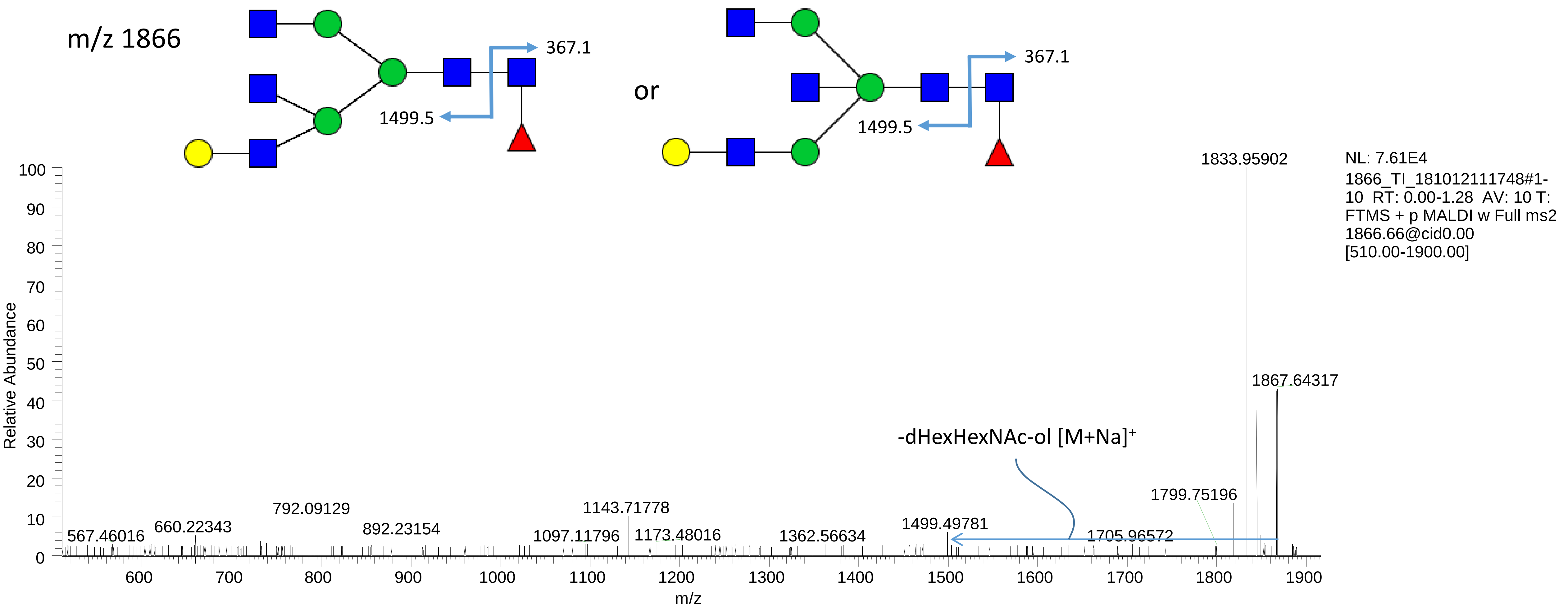
m/z 1866
367.1
1499.5
or
367.1
1499.5
1833.95902
NL: 7.61E4
1866_TI_181012111748#1-10 RT: 0.00-1.28 AV: 10 T: FTMS + p MALDI w Full ms2 1866.66@cid0.00 [510.00-1900.00]
1867.64317
-dHexHexNAc-ol [M+Na]+
1799.75196
1705.96572
1499.49781
1362.56634
1143.71778
1173.48016
1097.11796
892.23154
792.09129
660.22343
567.46016
Relative Abundance
m/z

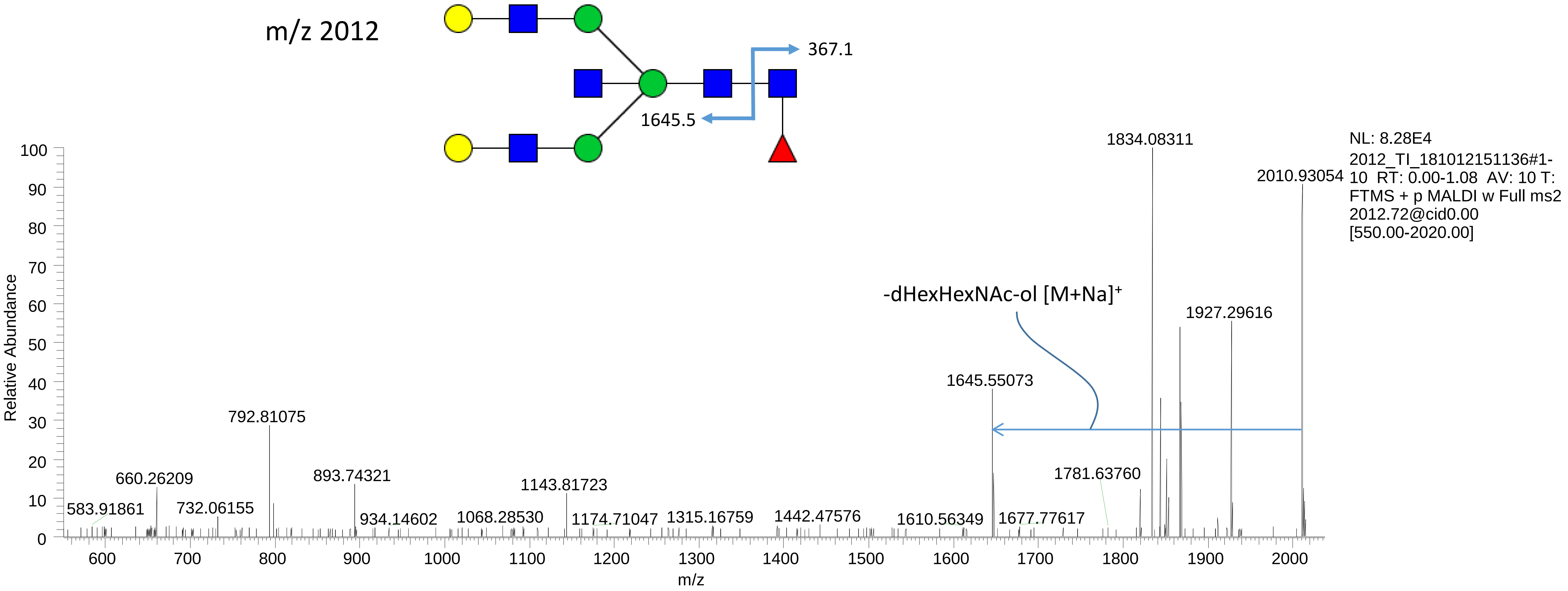
m/z 2012
367.1
1645.5
-dHexHexNAc-ol [M+Na]+
NL: 8.28E4
2012_TI_181012151136#1-10 RT: 0.00-1.08 AV: 10 T: FTMS + p MALDI w Full ms2 2012.72@cid0.00 [550.00-2020.00]
Relative Abundance
m/z
1834.08311
2010.93054
1927.29616
1645.55073
1781.63760
792.81075
660.26209
893.74321
1143.81723
583.91861
732.06155
934.14602
1068.28530
1174.71047
1315.16759
1442.47576
1610.56349
1677.77617
