## Supplemental Data 5_Total glycoforms detected for "On-tissue spatially-resolved glycoproteomics guided by N-glycan imaging reveal global dysregulation of canine glioma glycoproteomic landscape"

| m/z (RapiFlex) | ± Da | Modification or adduct | Glycan assignment | Glycan Class | Accurate mass (Orbitrap) | ± ppm |
|---|---|---|---|---|---|---|
| 1079.356 | 0.25 | | HexNAc2Hex3Fuc1 | | 1079.37567 | 0 |
| 1095.305 | 0.25 | | HexNAc2Hex4 | | 1095.36651 | 2.8 |
| 1257.378 | 0.25 | | HexNac2Hex5 | HighMannose | 1257.42541 | 1.6 |
| 1273.327 | 0.25 | 1257 + Ox | HexNac2Hex5 | HighMannose | 1273.39925 | |
| 1339.463 | 0.25 | | HexNAc4Hex3 | Complex | 1339.4791 | 2.3 |
| 1419.391 | 0.25 | | HexNac2Hex6 | HighMannose | 1419.47909 | 2.2 |
| 1435.4 | 0.25 | 1419 + Ox | HexNac2Hex6 | HighMannose | 1435.45279 | |
| 1485.467 | 0.25 | | HexNAc4Hex3Fuc | Complex | 1485.53682 | 1.4 |
| 1501.476 | 0.25 | | HexNac4Hex4 | Complex | 1501.51189 | 10 |
| 1508.791 | 0.25 | 1485 + Na | HexNAc4Hex3Fuc | Complex | 1508.81069 | |
| 1542.489 | 0.25 | | HexNAc5Hex3Fuc | Complex | 1542.56156 | 4 |
| 1581.523 | 0.25 | | HexNac2Hex7 | HighMannose | 1581.53148 | 1.9 |
| 1597.473 | 0.25 | 1581 + Ox | HexNac2Hex7 | HighMannose | 1597.50596 | |
| 1611.923 | 0.25 | Na+ K+ | HexNAc3Hex3FucNeuAc | Complex | Not detected | |
| 1647.479 | 0.25 | | HexNAc4Hex4Fuc | Complex | 1647.59005 | 1.9 |
| 1663.489 | 0.25 | | HexNAc4Hex5 | Complex | 1663.5849 | 1.8 |
| 1688.492 | 0.25 | | HexNAc5Hex3Fuc | Complex | 1688.6173 | 2.4 |
| 1704.538 | 0.25 | | HexNAc5Hex4 | Complex | 1704.59276 | 9 |
| 1743.476 | 0.25 | | HexNac2Hex8 | HighMannose | 1743.58383 | 1.8 |
| 1759.574 | 0.25 | 1743 + Ox | HexNac2Hex8 | HighMannose | 1759.56021 | |
| 1791.564 | 0.25 | | HexNAc4Hex4NeuAc | Complex | Not detected | |
| 1809.492 | 0.25 | | HexNAc4Hex5Fuc | Complex | 1809.64969 | 5.1 |
| 1825.442 | 0.25 | | HexNAc4Hex6 | Hybrid | 1825.62536 | 4.5 |
| 1850.565 | 0.25 | | HexNAc5Hex4Fuc | Complex | 1850.67171 | 2.8 |
| 1866.694 | 0.25 | | HexNAc5Hex5 | Complex | 1866.64874 | 6.6 |
| 1891.578 | 0.25 | | HexNAc6Hex3Fuc | Complex | 1891.6994 | 3.2 |
| 1905.549 | 0.25 | | HexNac2Hex9 | HighMannose | 1905.64036 | 2.1 |
| 1921.618 | 0.25 | 1905 + Ox | HexNac2Hex10 | HighMannose | 1921.61325 | |
| 1954.596 | 0.25 | | HexNAc4Hex5NeuAc1 | Complex | 1954.68023 | 1.6 |
| 1955.555 | 0.25 | | HexNAc4Hex5Fuc2 | Complex | Not detected | |
| 1976.542 | 0.25 | 1955 + Na+ | HexNAc4Hex5Fuc2 | Complex | 1976.71145 | |
| 1996.628 | 0.25 | | HexNAc5Hex4Fuc2 | Complex | 1996.72915 | 2.6 |
| 2012.638 | 0.25 | | HexNAc5Hex5Fuc | Complex | 2012.70866 | 5.1 |
| | | | HexNAc5Hex6 | Complex | 2028.71013 | 1.5 |
| 2053.651 | 0.25 | | HexNAc6Hex4Fuc | Complex | 2053.73851 | 3.5 |
| | | | HexNAc2Hex10 | HighMannose | 2067.68853 | 0.49 |
| | | | HexNAc4Hex5Fuc1NeuAc1 | Complex | 2100.73735 | 0.97 |
| 2101.679 | 0.25 | | HexNAc4Hex5Fuc3 | Complex | 2101.75548 | 0 |
| | | | HexNAc4Hex6NeuAc1 | Hybrid | 2116.71624 | 6.3 |
| 2117.688 | 0.25 | 2101 + Ox | HexNAc4Hex5Fuc3 | Complex | 2117.72578 | |
| 2133.698 | 0.25 | | HexNac4Hex7Fuc1 | Hybrid | 2133.74709 | 0.96 |
| 2158.641 | 0.25 | | HexNAc5Hex5Fuc2 | Complex | 2158.78202 | 2.4 |
| | | | HexNAc5Hex6Fuc | Complex | 2174.76899 | 0.94 |
| 2215.663 | 0.25 | | HexNAc6Hex5Fuc | Complex | 2215.7664 | 14 |
| 2245.597 | 0.25 | | HexNAc4Hex5NeuAc2 | Complex | 2245.77724 | 2.3 |
| 2261.653 | 0.25 | | HexNAc4Hex5NeuAcNeuGc | Complex | 2261.74712 | |
| 2267.589 | 0.25 | 2245 + Na+ | HexNAc4Hex5NeuAc2 | Complex | 2267.75387 | |
| 2283.598 | 0.25 | 2261 + Na+ | HexNAc4Hex5NeuAcNeuGc | Complex | 2283.72586 | |
| 2289.594 | 0.25 | 2245 + 2Na+ | HexNAc4Hex5NeuAc2 | Complex | 2289.73787 | |
| 2299.668 | 0.25 | Na+ | HexNAc4Hex5NeuGc2 | Complex | 2299.69253 | |
| 2305.604 | 0.25 | 2261 + 2Na+ | HexNAc4Hex5NeuAcNeuGc | Complex | 2305.80182 | |
| 2321 | 0.25 | 2291 + 2Na+ | HexNAc4Hex5NeuGc2 | Complex | 2321.71098 | |
| 2361.607 | 0.25 | | HexNAc6Hex5Fuc2 | Complex | 2361.85713 | 0.43 |
| | | | HexNAc4Hex5Fuc1NeuAc2 | Complex | 2391.83609 | 2.6 |
| 2406.277 | 0.25 | | HexNAc6Hex5Fuc2 | Complex | 2406.65863 | |
| | | | HexNAc4Hex6NeuAc2 | Complex | 2407.80509 | 8 |
| | | Na+ | HexNAc4Hex5Fuc1NeuAcNeuGc | Complex | 2429.78391 | |
| | | | HexNAc5Hex6Fuc1NeuAc1 | Complex | 2465.86279 | 1.6 |
| | | | HexNAc5Hex7NeuAc1 | | 2481.84185 | 7.8 |
| | | | HexNAc5Hex8Fuc1 | | 2498.87615 | 0 |
| | | | HexNAc5Hex9 | | 2514.87241 | 0 |
| | | | HexNAc6Hex7Fuc1 | | 2539.90269 | 0.4 |
| | | | HexNAc6Hex8 | | 2555.90380 | 1.6 |
| | | | HexNAc5Hex6Fuc2NeuAc1 | | 2611.91541 | 3.5 |
| | | | HexNAc5Hex7Fuc1NeuAc1 | | 2627.91403 | 1.9 |
| | | 2611 + Na+ | HexNAc5Hex6Fuc2NeuAc1 | | 2633.84988 | |
| | | | HexNAc5Hex8NeuAc1 | | 2643.89790 | 6.1 |
| | | | HexNAc5Hex9Fuc1 | | 2660.92800 | 0.38 |
| | | | HexNAc6Hex7Fuc2 | | 2685.94783 | 4.9 |
| | | | HexNAc6Hex9Fuc1 | | 2701.95058 | 2.2 |
| | | | HexNAc6Hex9 | | 2717.95486 | 1.2 |
| | | | HexNAc5Hex7Fuc2NeuAc1 | | 2773.95157 | 9.5 |
| | | | HexNAc5Hex8Fuc1NeuAc1 | | 2789.97431 | 0.36 |
| | | | HexNAc6Hex7Fuc1NeuAc1 | | 2830.98241 | 5.7 |
| | | | HexNAc6Hex7Fuc3 | | 2831.99565 | 8.6 |
| | | | HexNAc6Hex8Fuc2 | | 2847.99004 | 8.6 |
| | | | HexNAc6Hex9Fuc1 | | 2864.00465 | 1.8 |
| | | | HexNAc6Hex10 | | 2880.00599 | 0.7 |
| | | | HexNAc5Hex7Fuc3NeuAc1 | | 2920.02648 | 3.1 |
| | | | HexNAc6Hex8Fuc3 | | 2994.04947 | 7.8 |
