## Supplemental Data 6_Lectin Staining for "On-tissue spatially-resolved glycoproteomics guided by N-glycan imaging reveal global dysregulation of canine glioma glycoproteomic landscape"

**Supplementary Data 6. Lectin staining.** Using a Dako delimiting pen (Agilent Technologies, Santa Clara, CA), dams were created around tissue sections. The sections were then incubated in 1% BSA (w/v) in approximately 300 µL of PBS for 30 min at room temperature, then incubated in 10 µg/mL of SNA lectin for 2 h. They were then rinsed for 10 min three times with 1% BSA in PBS. The sections were then incubated in approximately 300 µL of DAPI for 20 min and rinsed with PBS for 5 min. Finally, two drops of Vectashield fluorescence mounting medium (Dako, Agilent Technologies) was added and the sections were cover-slipped and sealed with nail polish. Confocal images were obtained using a fluorescence microscope (Leica Biosystems). Adjacent tissue sections incubated in 1% BSA in PBS served as controls. Zeiss LSM700 confocal microscope connected to a Zeiss Axiovert 200 M with an EC Plan-Neofluar 40x/1.30 numerical aperture oil immersion objective (Carl Zeiss AG, Oberkochen, Germany). Processing of the images was performed using Zen software and applied on the entire images as well as on controls. The presented pictures are representative of independent triplicates.

# VH14-0622

Glioblastoma (Grade IV WHO)

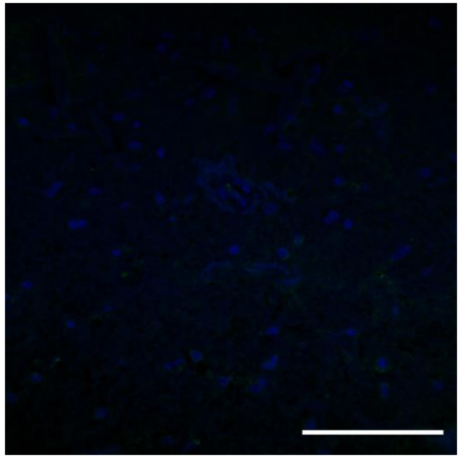

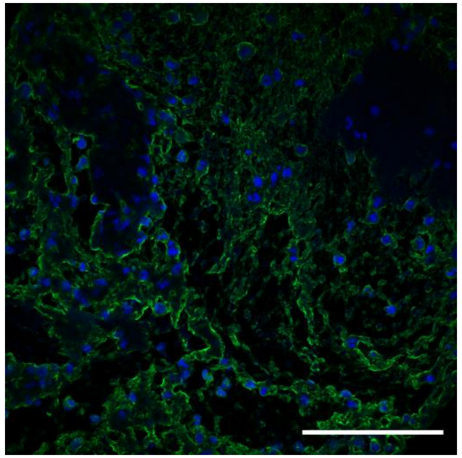

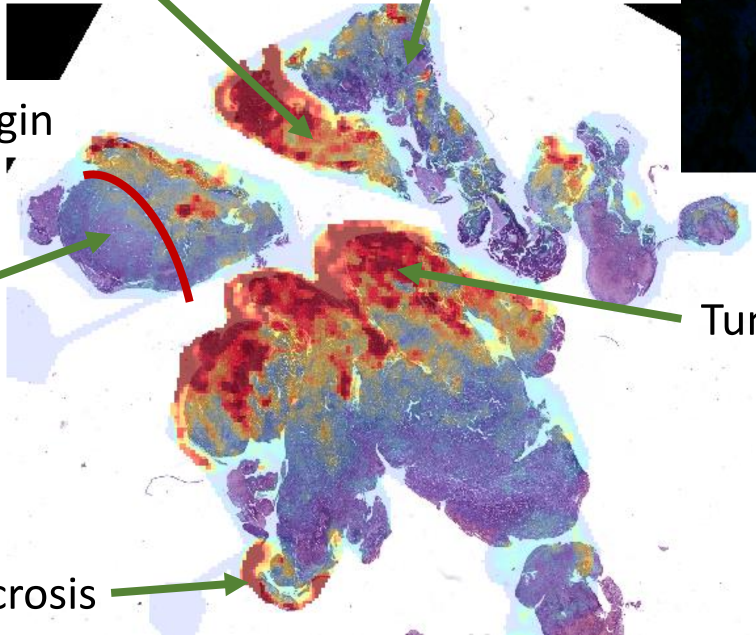


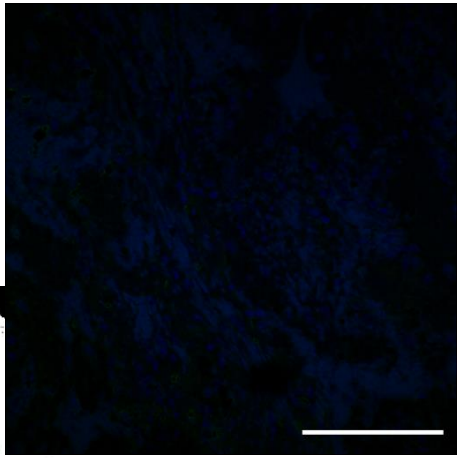

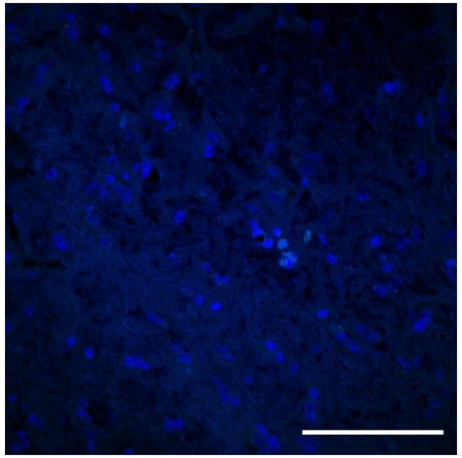

Control: separate section

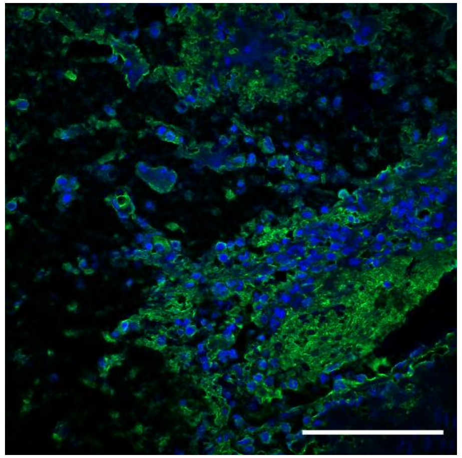

Green=positive for SNA lectin staining
Blue=DAPI

# VH15-1139A

Glioblastoma (Grade IV WHO)

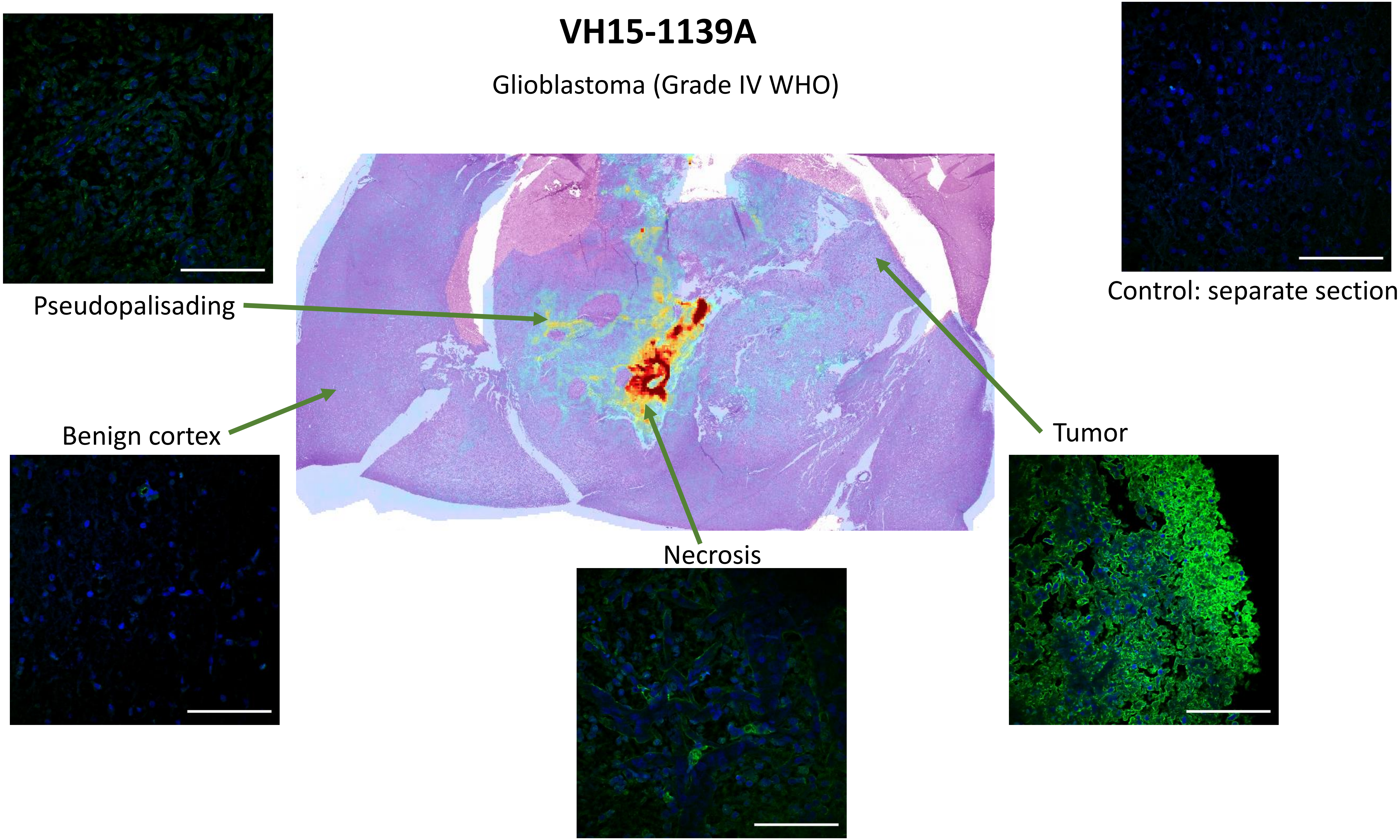

# VH15-1139D

Glioblastoma (Grade IV WHO)

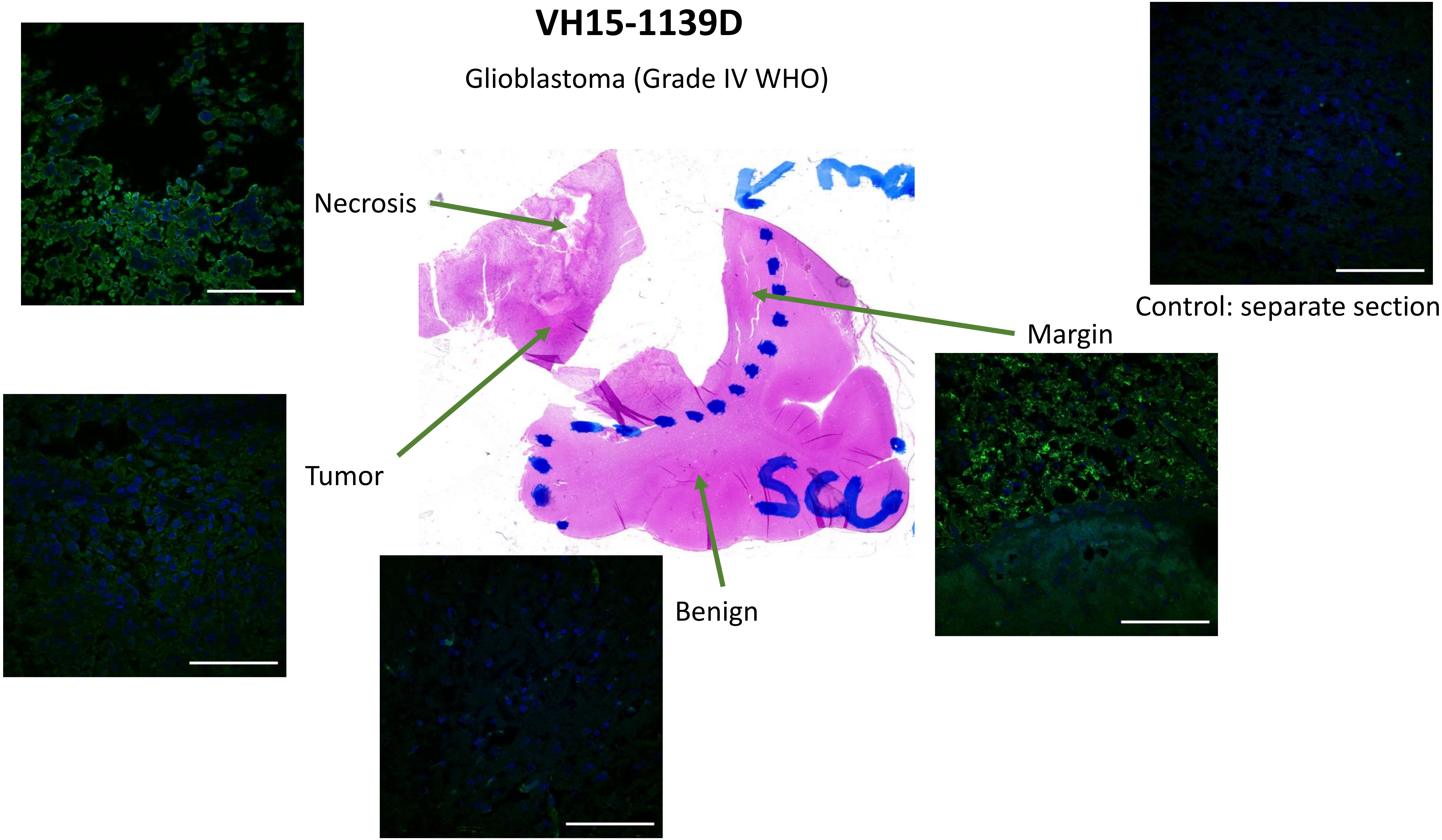

# VH15-3520A

Anaplastic Oligodendroglioma (Grade III WHO)

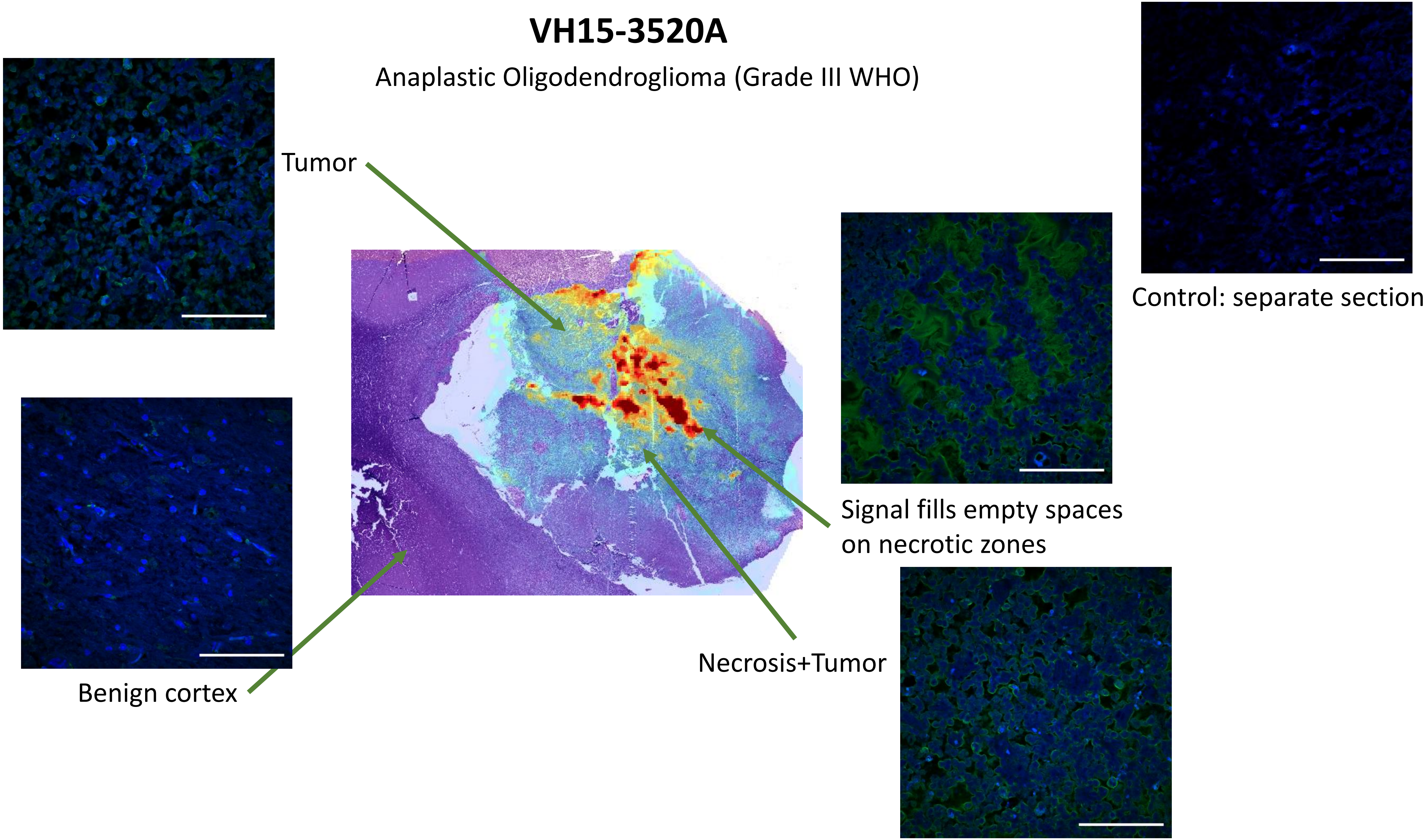

# VH16-0703A

Anaplastic Oligodendroglioma (Grade III WHO)

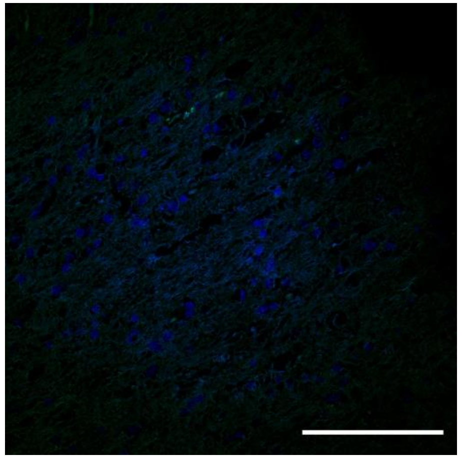

Control: separate section

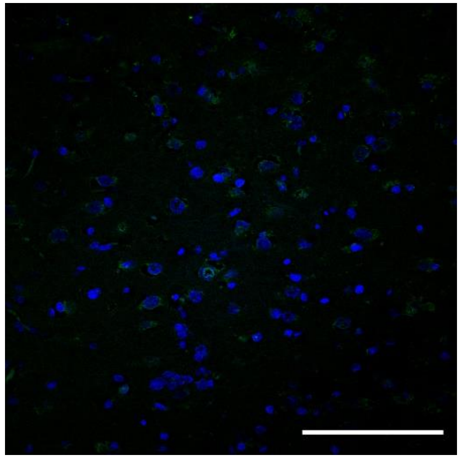

Benign – No signal on Cortex

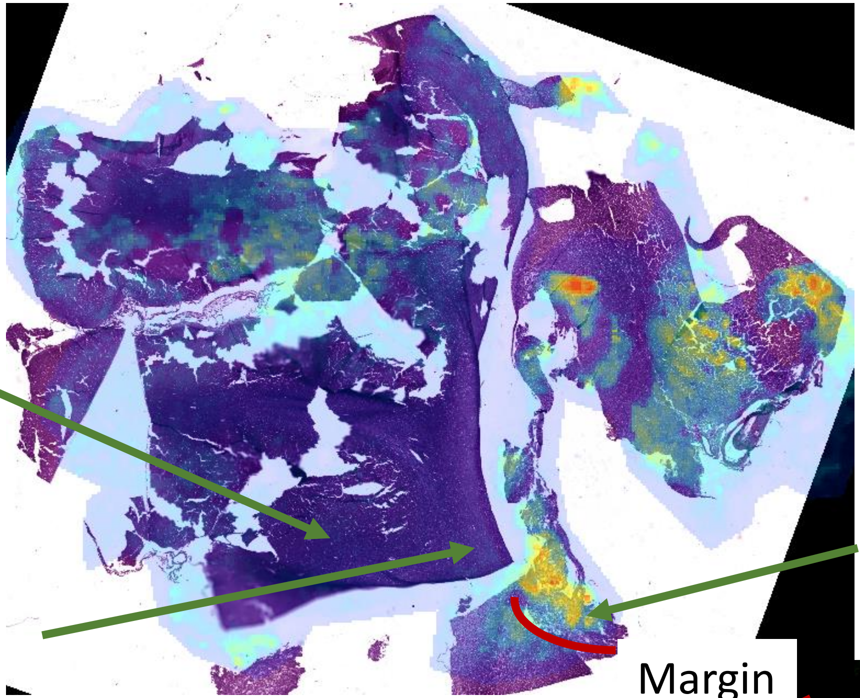

Necrosis

Margin

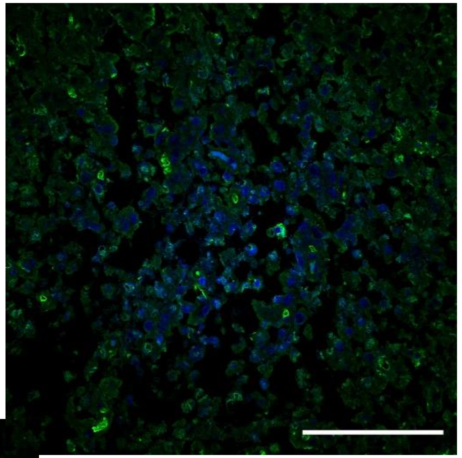

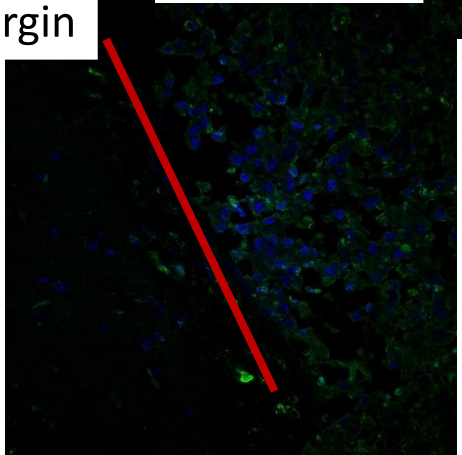

Benign – Intense signal on corpus callosum

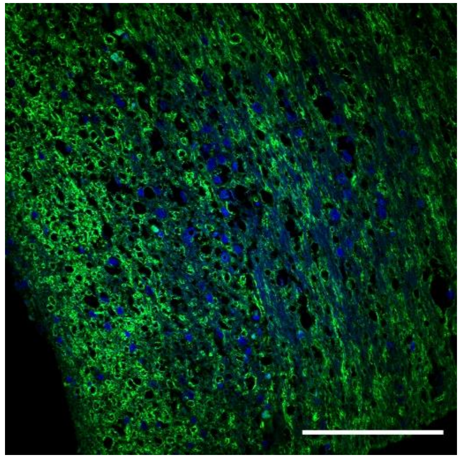

# VH16-0703B

Anaplastic Oligodendroglioma (Grade III WHO)

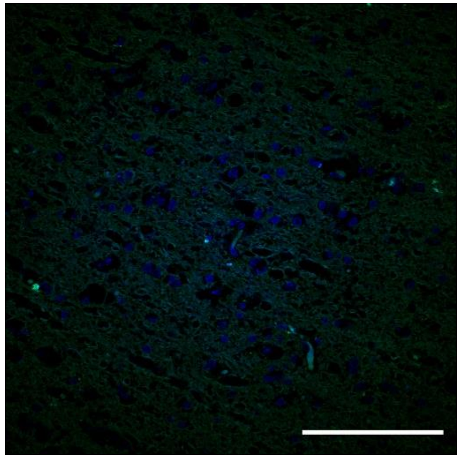

Control: separate section

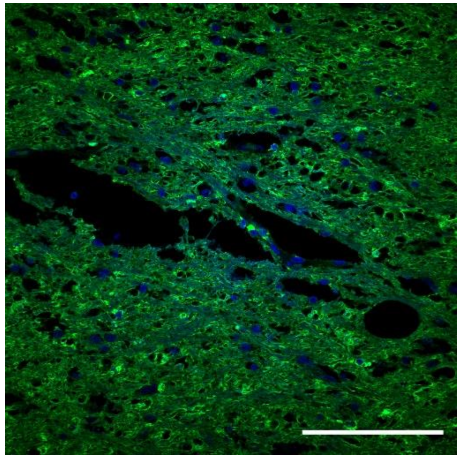

Benign – Intense signal on corpus callosum

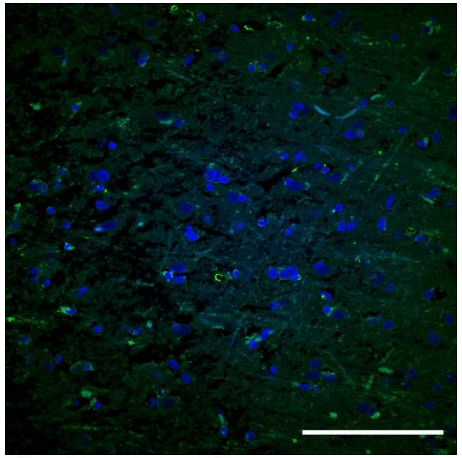

Benign – No signal on Cortex

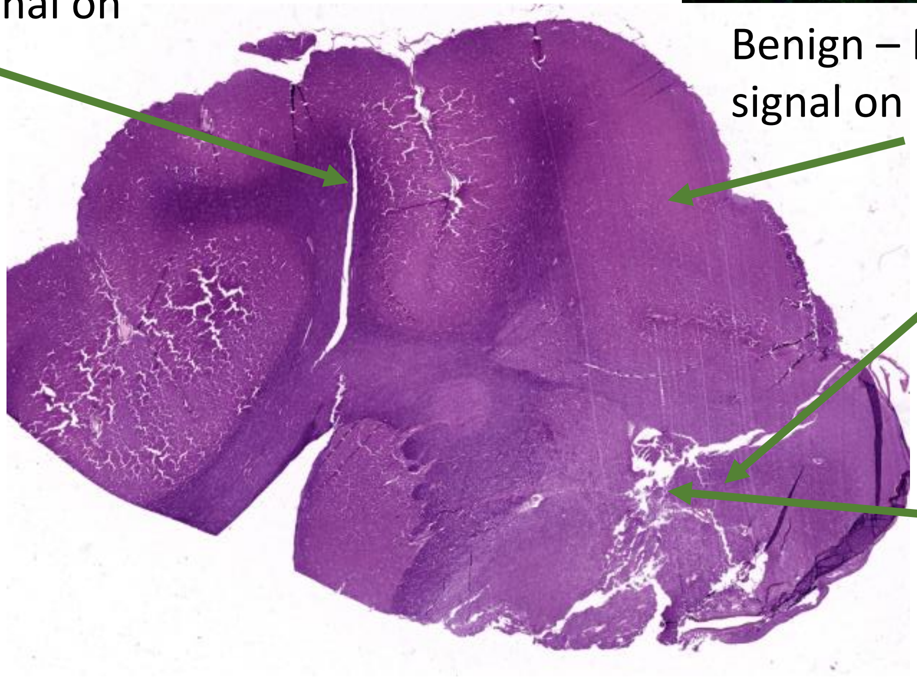

Tumor

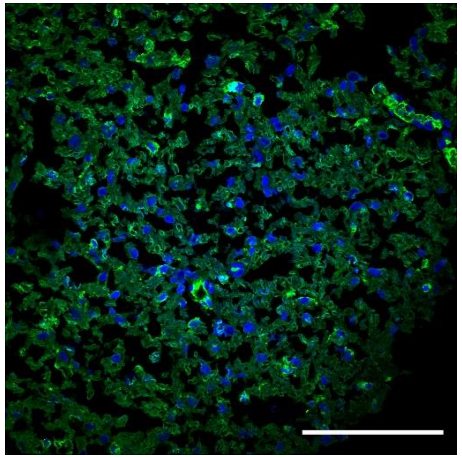

Necrosis

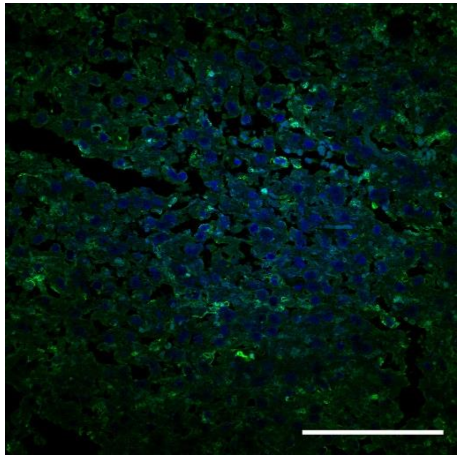

# VH13-0935

Anaplastic Oligodendroglioma (Grade III WHO)

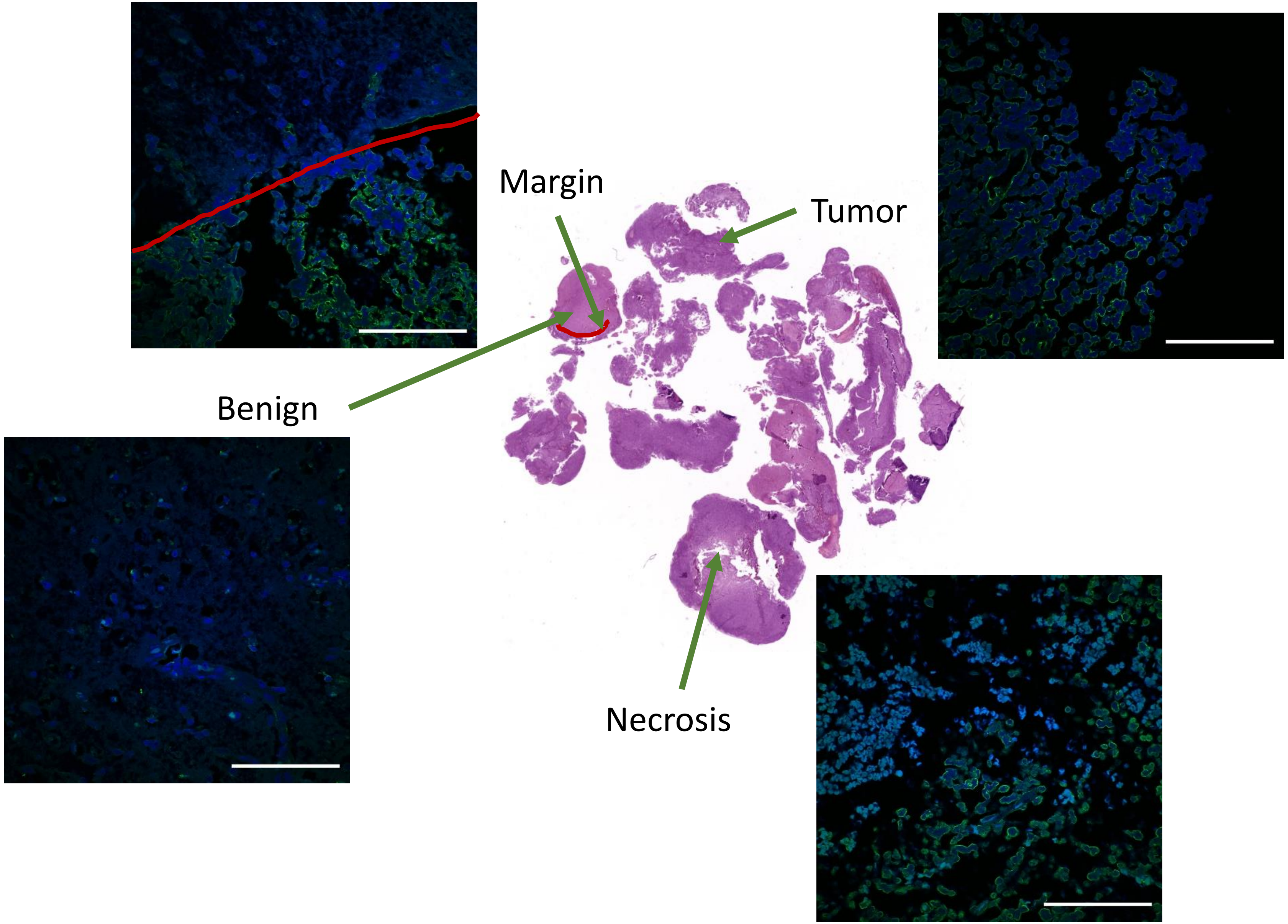


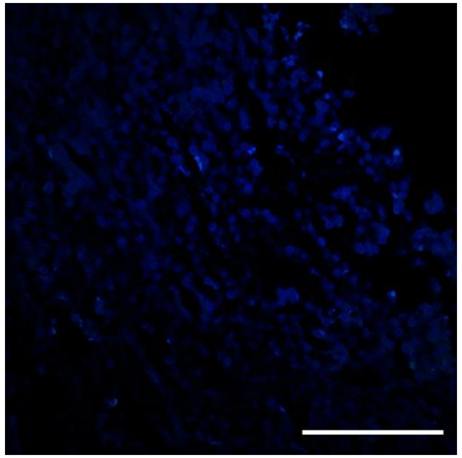

Control: separate section

# VH16-0440C

Anaplastic Oligodendroglioma (Grade III WHO)

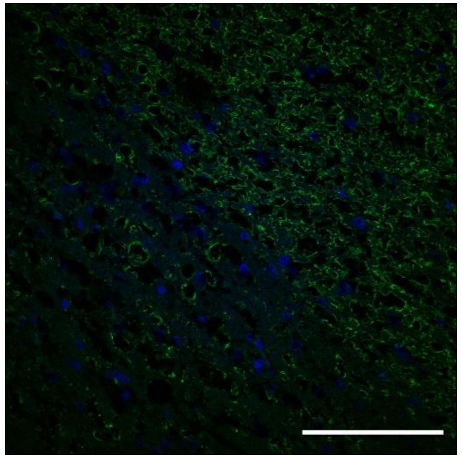

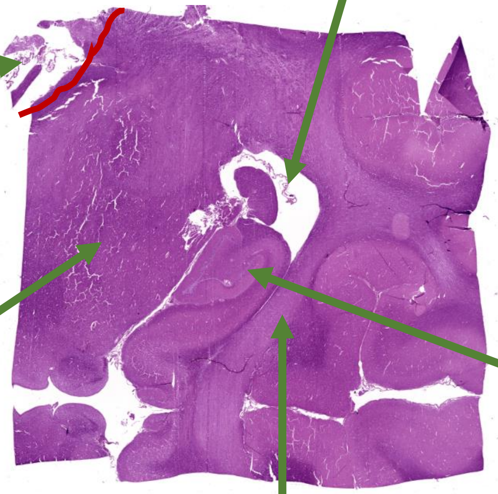


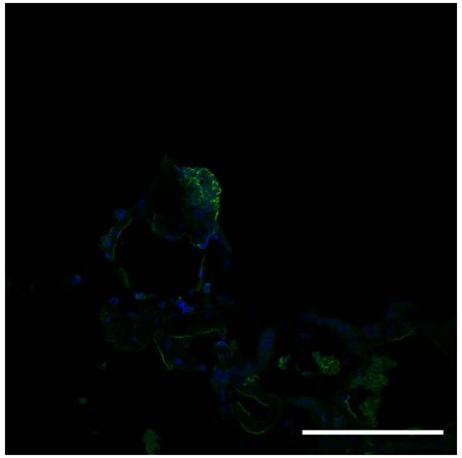

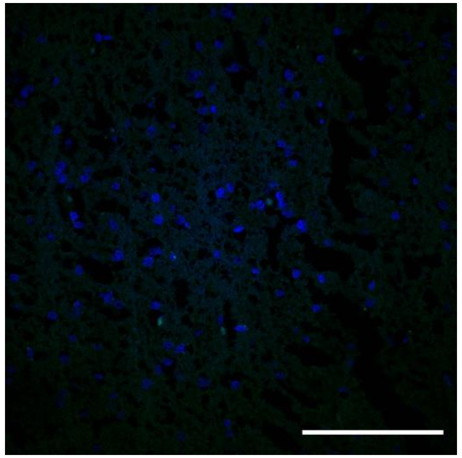

Control: separate section

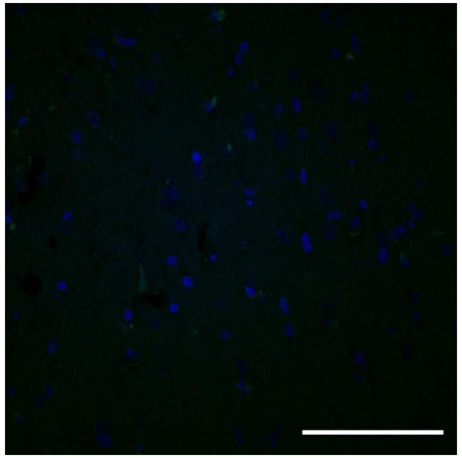

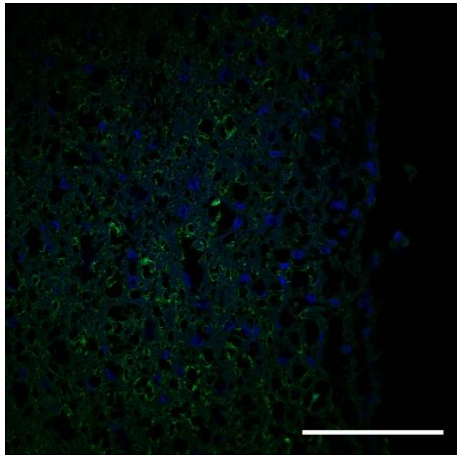

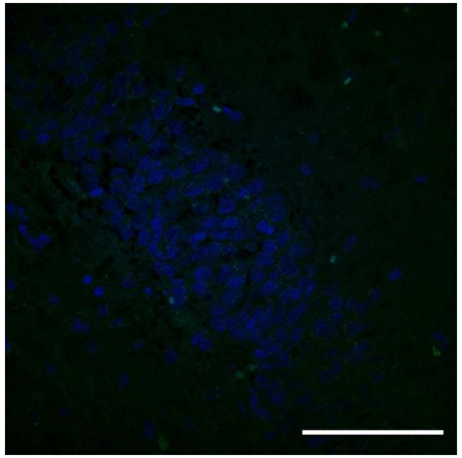

# VH16-0440D

Anaplastic Oligodendroglioma (Grade III WHO)

No visible tumor/necrotic zone

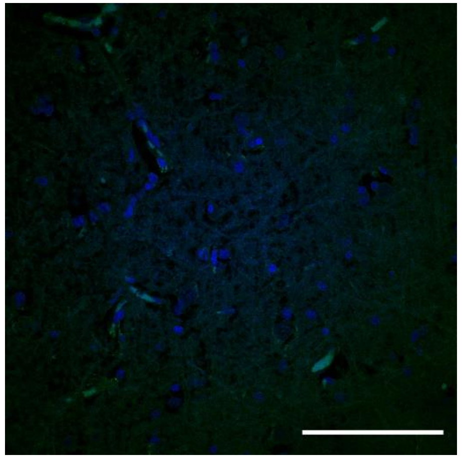

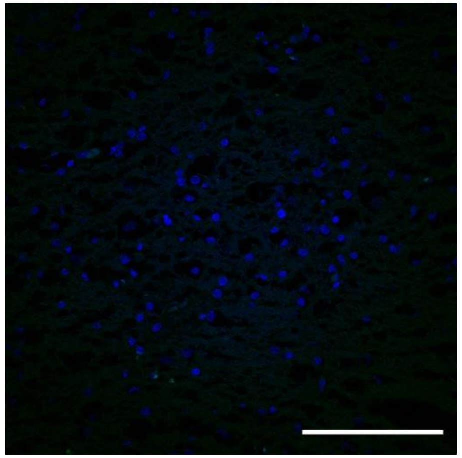

Control: separate section

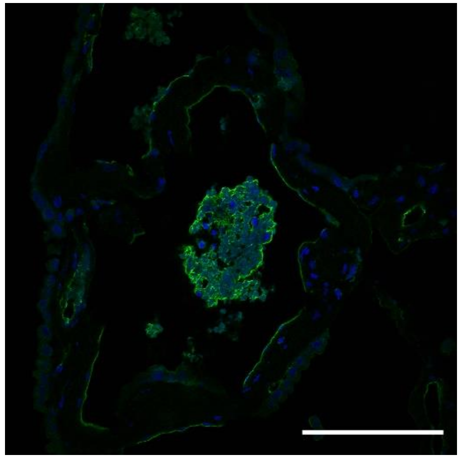

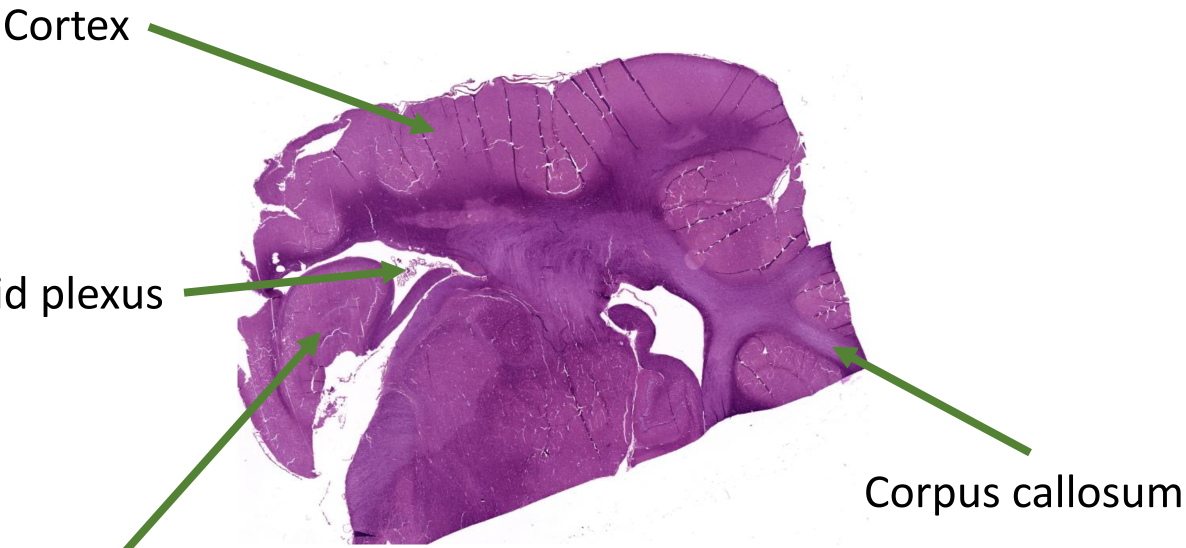


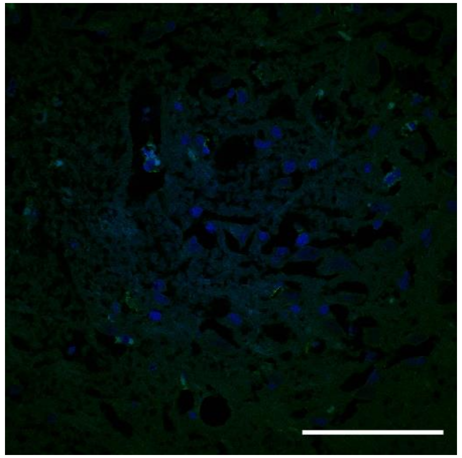

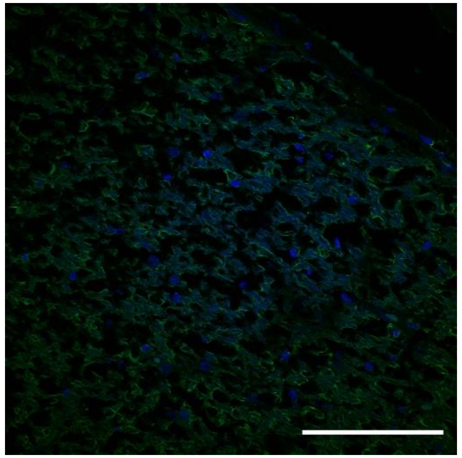
